# A TrkB and TrkC partial agonist restores deficits in synaptic function and promotes activity-dependent synaptic and microglial transcriptomic changes in a late-stage Alzheimer’s mouse model

**DOI:** 10.1101/2023.09.18.558138

**Authors:** Amira Latif-Hernandez, Tao Yang, Robert Raymond-Butler, Patricia Moran Losada, Paras Minhas, Halle White, Kevin C. Tran, Harry Liu, Danielle A. Simmons, Vanessa Langness, Katrin Andreasson, Tony Wyss-Coray, Frank M. Longo

## Abstract

**Introduction:** TrkB and TrkC receptor signaling promotes synaptic plasticity and interacts with pathways affected by amyloid-β (Aβ)-toxicity. Upregulating TrkB/C signaling could reduce Alzheimer’s disease (AD)-related degenerative signaling, memory loss, and synaptic dysfunction.

**Methods:** PTX-BD10-2 (BD10-2), a small molecule TrkB/C receptor partial agonist, was orally administered to aged London/Swedish-APP mutant mice (APP^L/S^) and wild-type controls (WT). Effects on memory and hippocampal long-term potentiation (LTP) were assessed using electrophysiology, behavioral studies, immunoblotting, immunofluorescence staining, and RNA-sequencing.

**Results:** Memory and LTP deficits in APP^L/S^ mice were attenuated by treatment with BD10-2. BD10-2 prevented aberrant AKT, CaMKII, and GLUA1 phosphorylation, and enhanced activity-dependent recruitment of synaptic proteins. BD10-2 also had potentially favorable effects on LTP-dependent complement pathway and synaptic gene transcription.

**Conclusions:** BD10-2 prevented APP^L/S^/Aβ-associated memory and LTP deficits, reduced abnormalities in synapse-related signaling and activity-dependent transcription of synaptic genes, and bolstered transcriptional changes associated with microglial immune response.

## 1. INTRODUCTION

Synaptic dysfunction and eventual loss of synapses are fundamental contributors to cognitive deficits in Alzheimer’s disease (AD) [1, 2]. Other key neuropathological features of AD include accumulation of toxic species of amyloid β (Aβ) and tau proteins. Pathological Aβ and tau attenuate hippocampal long-term potentiation (LTP) in AD mouse models [3–5]. LTP is a form of synaptic plasticity involved in memory and learning. LTP impairments induced by Aβ and tau can be explained, in part, by disrupted neurotrophin signaling [6–11]. The neurotrophin, brain-derived neurotrophic factor (BDNF), is released in an activity-dependant manner in the hippocampus. BDNF binds to and activates tropomyosin related kinase B (TrkB) receptors. Upon activation, TrkB translocates to the postsynaptic terminal [12] and initiates signaling cascades including the phosphatidylinositol-3 kinase (PI3K)/AKT and phospholipase Cy-1 (PLCγ)/protein kinase C (PKC) pathways, which mediate LTP maintenance and support synaptic transmission and membrane excitability [13]. Synaptic plasticity is also facilitated by the neurotrophin NT-3, and signaling via its receptor, TrkC [14, 15]. Deficiencies in BDNF/TrkB and NT-3/TrkC signaling have been observed in the brains of AD patients and mouse models, and have been associated with Aβ and tau accumulation, synapse loss, and cognitive decline [16–18]. TrkB/TrkC signaling pathways largely intersect molecular pathways essential to Aβ and tau pathology and contribute to synaptic plasticity. Thus, activating TrkB/TrkC signaling may counteract AD-related degenerative signaling and restore LTP.

This study examined whether promoting TrkB and TrkC signaling with a small molecule ligand can reduce synaptic plasticity deficits in the hAPP^Lond/Swe^ (APP^L/S^) mouse model of AD. Previously, we identified a small molecule, LM22B-10, that activates TrkB and TrkC and their downstream signaling pathways, including PI3K/AKT, MAPK/ERK and PLCγ/PKC while preventing dendritic spine loss in aged mice [19]. A derivative of LM22B-10, designated as PTX-BD10-2 (BD10-2), was developed which has improved blood brain barrier penetration following oral administration and reverses cholinergic neuron atrophy associated with pathological tau (AT8 antibody) in late-stage APP^L/S^ mice [20]. In the present investigation, BD10-2 was orally administered to APP^L/S^ mice with memory deficits and late-stage amyloid pathology [21, 22]. This therapeutic strategy was adopted to increase clinical relevance as therapeutic trials are generally initiated when key elements of pathology are already established. We found that BD10-2 significantly reduced memory and LTP impairments, as well as synaptic- and neurotrophin-related signaling deficits in APP^L/S^ mice. These beneficial effects may be explained in part by BD10-2’s ability to normalize alterations in activity-dependent transcription of synaptic genes that were observed in vehicle treated APP^L/S^ mice. Additionally, we identified upregulation of genes in pathways associated with microglia and immune responses in APP^L/S^ mice that were further upregulated in APP^L/S^ mice treated with BD10-2 suggesting that BD10-2 may enhance one or more elements of the existing immune response to AD-related pathology.

## 2. METHODS

### 2.1 Reagents

LM22B-10 (2-[[4-[[4-[Bis-(2-hydroxy-ethyl)-amino]-phenyl]-(4-chloro-phenyl)-methyl]-phenyl]-(2-hydroxy-ethyl)-amino]-ethanol) and PTX-BD10-2 (BD10-2) (Bis-{4-[bis-(2-methoxy-ethyl)-amino]-phenyl}-ethane) were custom manufactured for our laboratory by Ricerca Biosciences LLC (Concord, OH). The compounds were characterized by high-performance liquid chromatography (HPLC), and liquid chromatography/mass spectrometry (LC-MS) and had a purity of ≥ 97%. Recombinant BDNF was purchased from PeproTech (Rocky Hill, NJ). Other reagents were purchased from Sigma-Aldrich Corp (St. Louis, MO), unless otherwise stated.

### 2.2 NIH-3T3 cell cultures and cell survival assays

Mouse NIH-3T3 cells expressing TrkA (NIH-3T3-TrkA) and p75 (NIH-3T3-p75) were provided by Dr. William Mobley (University of California San Diego), and NIH-3T3 cells expressing TrkB (NIH-3T3-TrkB) or TrkC (NIH-3T3-TrkC) were provided by Dr. David Kaplan (University of Toronto). Cells were propagated in DMEM supplemented with 10% FBS (Invitrogen) and 200-400 µg/ml Geneticin. Cells were seeded onto 24-well plates (30,000 cells/well) and cultured in medium consisting of 50% PBS and 50% DMEM without supplements. Following exposure to BDNF (20 ng/ml, 0.7 nM) or 10-1000 nM LM22B-10 or BD10-2 for 72-96 h, cells were suspended in 50 µl lysis buffer, transferred to white, opaque 96-well culture plates and survival was measured using the ViaLight Assay (Lonza Group Ltd., Rockland, ME).

### 2.3 Primary neuronal cultures and cell survival assays

Hippocampal neuron cultures were prepared from embryonic day 16 (E16) CF1 mouse fetuses, as described previously [23]. Under the low-density conditions used in the present studies, neuronal survival is dependent, in part, on addition of exogenous neurotrophins [24, 25]. LM22B-10 and BD10-2 were dissolved in water with 2-3% of 1N HCl at a stock concentration of 10 mM prior to dilution (1:10,000) in culture medium. After 48 h exposure to BDNF, LM22B-10, or BD10-2, cells were immunostained for the neuron-specific marker, β-tubulin III (Tuj1), and cell survival was quantified by counting immunopositive, morphologically intact cell bodies [25, 26].

### 2.4 Animals

All animal use procedures complied with the National Institutes of Health Guide for the Care and Use of Laboratory Animals and all protocols were approved by the Institutional Animal Care and Use Committee at Stanford University. These experiments used Thy1-hAPP^Lond/Swe^ [Line 41] mice (APP^L/S^) which express human APP751 containing the London (V7171) and Swedish (K670M/N671L) mutations under the control of the Thy-1 promoter [21]. Mice were bred in our laboratory and maintained on a C57BL/6 background. Mice were group housed in mixed genotypes. All mice received cotton nestlets and paper tubes. Water and food were freely available. Genotyping using tail DNA was performed by TransnetYX Inc.

### 2.5 Study Design for BD10-2 Treatment of APP^L/S^ mice

To examine the acute effects of BD10-2 on LTP, hippocampal slices were prepared from a group of male 16-month-old wild-type (WT; n = 6 mice) and APP^L/S^ mice (n = 6 mice). BD10-2 was dissolved by agitation and sonication at 37 °C in 25% hydroxypropyl-β-cyclodextrin (HPCD; Aldrich) in de-ionized water as the vehicle. Hippocampal slices were exposed to 1000 nM BD10-2 or vehicle via bath application. A separate group of male (WT; n = 34) and APP^L/S^ (n = 44) mice were administered vehicle or BD10-2 [50 mg/kg, 10 ml/kg, once daily by oral gavage (O.G.) five days/week for 3 months starting at 13 months of age. A 2 × 2 study design was used with the following groups: WT-Veh (n = 17 mice), WT-BD10-2 (n = 17 mice), APP-Veh (n = 22 mice) and APP-BD10-2 (n = 22 mice). BD10-2 was administered to late-symptomatic APP^L/S^ mice when extracellular Aβ deposits (present at age 3-5 months), tau pathology (present at age 6-8 months), and memory deficits (present at age 6-9 months) are well-established [20–22, 27–29]. After ∼2 months of treatment, memory-related performance in WT-Veh (n = 10 mice), WT-BD10-2 (n = 10 mice), APP-Veh (n = 9 mice) and APP-BD10-2 (n = 12 mice) mice was assessed using the novel object recognition (NOR), novel object displacement (NOD), and Barnes maze tests. Behavioral testing was performed daily, 24 h after the last BD10-2 or vehicle dose for ∼1.5 months. A separate cohort of BD10-2 or vehicle treated mice [WT-Veh (n = 7 mice), WT-BD10-2 (n = 7 mice), APP-Veh (n = 13 mice) and APP-BD10-2 (n = 10 mice)] were euthanized at 16 months of age and hippocampal slices were prepared for electrophysiology recording 24 h after the last BD10-2 or vehicle administration to measure LTP.

### 2.6 Novel Object Recognition and Displacement

The novel object recognition (NOR) and novel object displacement (NOD) tasks are based on the ability of mice to show preference for novel objects versus familiar objects when allowed to explore freely [30]. Mice were individually habituated to an open arena (50 cm × 50 cm, dim light, 24 °C) 5 min prior to the training session with an inter-training interval (ITI) of 3 min. During the training session, two identical objects were placed into the arena and the animals were allowed to explore for 10 min. During the testing session for NOD, after a delay of 24 h, the animals were placed back into the same arena in which one of the objects used during training was moved to a new location in the arena. The mice were allowed to explore freely for 10 min. During the testing for NOR, the familiar object that remained in the same location on testing day 1 was replaced by a novel object of similar dimensions but with a different shape/color. The mice were allowed to explore freely for 10 min. Digital video tracking (using Kinovea software) of body movements and nose position was used to quantify the exploratory activity around the objects (2 cm zone around the objects). Exploration behavior was assessed by establishing the discrimination index (DI, as a percentage), i.e. the ratio of the time spent exploring the novel object over the time spent exploring the two objects. A DI of ≥ 50% is therefore characteristic of training; significantly increased DI is characteristic of recognition. To evaluate memory, comparisons were made for each genotype/treatment group between the testing sessions (24 h) and the training sessions (0 h). Behavioral experiments were performed by experimenters blinded to genotype and treatment.

### 2.7 Barnes Maze

The Barnes maze is a spatial memory task in which a mouse must learn a target location (hole) to escape a brightly lit platform using distal cues. The experimental protocol was adopted from [31] with minor modifications. The maze consisted of a circular, white PVC slab (8 mm thick, 91.4 cm diameter) with 20 holes (7.62 cm diameter) along the perimeter 2.54 cm from the edge and mounted on a rotating stool 76.2 cm above the ground. Beneath the escape hole, a platform and ramp was located 5.08 cm below the surface of the maze leading to a mouse cage with darkened walls to reduce light. The maze was situated in the center of a room with two 120 W lights facing toward it to provide an aversive stimulus. Eight simple colored-paper shapes (squares, rectangles, and circles) were mounted on the walls of the room as visual cues. After testing each mouse, the maze was cleaned with 70% ethanol and rotated clockwise to avoid intra-maze odor or visual cues. All sessions were recorded using a JVC Everio HD camcorder GZ-E200 and analyzed with Kinovea video tracking software. The animals interacted with the Barnes maze in three phases: habituation (day 1), training (days 2-3), and probe (day 4). Before starting each session, mice were acclimated to the testing room for 1 h. Then all mice from one cage (n = 4-5) were placed in individual holding cages where they remained until the end of their testing sessions each day. On habituation day, the mice were placed in the center of the maze within a vertically oriented black PVC pipe (10.2 cm diameter, 17.8 cm height) for 15 s and guided slowly to the hole that led to the escape cage over the course of 10-15 s. The mice were given 3 min to independently enter the target hole, and if they did not, they were nudged with the PVC pipe to enter. The 120 W lights were then shut off and mice were allowed to rest in the escape cage for 2 min. The training phase occurred 24 h after the habituation phase and was split across 2 days (days 2 and 3), with 3 trials on the first day and 2 trials on the second day. During each trial, the mice were placed in the center of the maze within the PVC pipe for 15 s and after allowed 3 min to explore the maze. If mice found and entered the target hole before 3 min passed, the lights were shut off and the training trial ended. Mice were allowed to rest in the escape cage for 2 min. If at the end of the 3 min the mice had not entered the target hole, they were nudged with the PVC pipe. A total of 5 trials were conducted. During each trial, latency (time) to enter the target hole as well as distance traveled were recorded. The probe phase occurred 24 h after the training phase and was conducted on the last day (day 4). Mice were placed in the center of the maze within the PVC pipe for 15 s and after allowed 3 min to explore the maze. The probe session ended whenever the mouse entered the target hole or if 3 min had passed. During the probe phase, measures of time spent per quadrant, latency to enter the target hole, and distance traveled were recorded. Behavioral experiments were performed by experimenters blinded to genotype and treatment.

### 2.8 Electrophysiology Recordings

Electrophysiological recordings were performed in hippocampal slices as previously described [32]. WT-Veh, WT-BD10-2, APP-Veh, and APP-BD10-2 slices were randomized during the experiment. For experiments using chronically treated mice, brains were harvested, and hippocampal slices were prepared 24 h after the final dose of BD10-2 or Vehicle (Veh). Animals were euthanized by cervical dislocation, the brain was removed, and the right hippocampus was rapidly dissected and placed into ice-cold (4 °C) artificial cerebrospinal fluid (ACSF) and saturated with carbogen (95% O_2_/5% CO_2_). The composition of the ACSF was as follows (in mM): 124 NaCl, 4.9 KCl, 24.6 NaHCO3, 1.20 KH2PO4, 2.0 CaCl2, 2.0 MgSO4, and 10.0 glucose, adjusted to a pH of 7.4. Transverse hippocampal slices were then prepared at a 350 μm thickness, from the dorsal region of the hippocampus using the McIlwain tissue chopper (Stoelting, Wood Dale, IL). These slices were transferred to a recovery chamber and maintained in oxygenated ACSF at room temperature for a minimum of 1.5 h. Slices were then placed into a submerged-type chamber at 32 °C where they were continuously perfused with ACSF at a flow-rate of 1.5 ml/min. Slices were delicately positioned onto an R6501A multi-electrode array (Alpha MED Scientific, Osaka, Japan) arrayed in an 8 × 2 matrix with electrodes spaced an interpolar distance of 150 μm and each matrix measured at 50 μm × 50 μm. Following a 30 min incubation period, field excitatory post synaptic potentials (fEPSPs) were recorded by stimulating downstream electrodes in the CA1 and CA3 regions along the Schaffer collateral pathway. The MED64 System (AlphaMED Sciences, Osaka, Japan) was used to acquire signals. The time course of the fEPSP was determined using the descending slope function for all the experiments. Input/output curves were generated by gradually increasing stimulus currents in the pathway, ranging from 10 μA to 90 μA (in 5-μA increments), while monitoring evoked responses. Once the input/output curves were established, the stimulation strength was adjusted to maintain the fEPSP slope at 35% of its maximal value throughout the experiment. During the baseline recording, a single response was evoked at a 30 s interval for at least 20 min. To induce a strong form of LTP, three episodes of theta-burst stimulation (TBS) were used, with each TBS consisting of 10 bursts of 4 stimuli at 100 Hz separated by 200 ms (double pulse-width). This was followed by a recording of evoked responses beginning 1 min after LTP induction and continuing every 30 s for 120 min after TBS application which marked the end of the experiment. The mean baseline fEPSP value was calculated and LTP was measured as the percentage change from baseline after TBS. Paired-pulse ratio (PPR) experiments were conducted, as previously described [7, 33] PPR responses to two impulses given at an interval of 10, 20, 50, 100, 200, or 500 ms were recorded. During baseline recording, 3 single stimuli (0.1 ms pulse width; 10 s intervals) were measured every 5 min and averaged for the 60 min fEPSP values. For experiments examining acute effects of BD10-2, the TrkB/C ligand was dissolved in water with 2-3% of 1N HCl at a stock concentration of 10 mM prior to dilution to a final concentration of 100 nM in the ACSF bath. Compound and vehicle solutions were added to oxygenated ACSF from 20 min before and until 130 min after TBS (see figures for details).

### 2.9 Protein extraction and Western blot analysis

Hippocampal slices with or without stimulation (TBS) were lysed in RIPA lysis buffer (150 mM NaCl, 50 mM Tris, pH 7.4, 1 mM EDTA, 1% Triton X-100 or 1% NP40, 10% glycerol, 1 mM PMSF, Na3VO4 and protease inhibitor cocktail). Lysates were mixed with 4X NuPage LDS loading buffer containing DTT. They were loaded (20-40 µg protein / lane) onto a NuPAGE 4-12% Bis-Tris gradient gel then transferred to PVDF membranes using 100 V for 1.5 h. Membranes were incubated in blocking buffer for 1 h at room temperature and then probed with primary antibodies. Antibodies consisted of: mouse monoclonal anti-phospho-ERK^T202/Y204^ (1:2000, 9106), rabbit polyclonal anti-ERK (1:2000, 9101), mouse monoclonal anti-phospho-AKT^S473^ (1:2000, 4051), rabbit polyclonal anti-AKT (1:2000, 9272), rabbit polyclonal anti-phospho-CaMKII^Thr286^ (1:2000, 12716), rabbit polyclonal CaMKII (1:2000, 3362), rabbit monoclonal anti-phopho-GluA1^Ser831^ (1:2000, 75574) and rabbit monoclonal GluA1 (1:2000, 13185) purchased from Cell Signaling Technology, Inc., (Danvers, MA); mouse anti-monoclonal PSD95 (1:3000, MAB1598) and synaptophysin (1:10000, MAB329-C) (Burlington, MA), mouse monoclonal anti-actin (1:10000, A5441) from Millipore Sigma. Primary antibody incubations were conducted overnight at 4 °C followed by incubation with the appropriate HRP conjugated secondary antibody for 1 h at room temperature. The immunobands were developed by incubating membranes in ECL developing solutions mixed in equal volumes for 1 min followed by imaging with Kodak film.

### 2.10 Double immunofluorescence staining of hippocampal slices

After electrophysiology experiments, hippocampal slices were post-fixed in 4% paraformaldehyde in phosphate-buffered saline (PBS) for 24 h and transferred to 30% sucrose (Sigma-Aldrich). Slices were sectioned coronally at 20 μm and stored at −20 °C in cryoprotectant (20% glycerol, 30% ethylene glycol in phosphate buffer) until immunostaining. Slices were washed twice for 5 min in tris-buffered saline (Tris-BS; pH 7.6), pretreated with 0.4% TritonX-100 in Tris-BS, followed by a 1 h incubation at room temperature in blocking solution (3% normal donkey serum, 0.4% TritonX-100, and 3% bovine serum in Tris-BS). Next, slices were incubated with primary antibodies for drebrin (1:1000, Enzo, Cat#ADI-NBA-110) and GLUA1 (AMPA receptor 1, GLUA1, 1:400, Cell Signaling Cat#13185) in blocking buffer overnight at 4 °C. They were then washed 3 times for 5 min and then incubated with secondary antibody Cy3-Donkey anti-rabbit (1:400) and FITC-Donkey anti-mouse (1:400) in Tris-BS for 4 h. Sections were mounted onto slides, cover-slipped with Prolog Anti-Fade solution with DAPI and imaged at least 48 h later.

### 2.11 Fluorescence Microscopy and Quantitative Image Analysis

Fluorescent images of mouse hippocampal slices immunostained for GluA1 and drebrin, were obtained using a Leica DM550 confocal microscope. A 63x objective was used to obtain z-stack images (0.7 μm step size; 40 steps) of the CA1 region of the hippocampus containing the most synapses near the pyramidal cells; 3 images were taken per slice. The laser power, gain, and offset settings were adjusted using LAS-X viewer software (Leica) to optimize signal capture; identical settings were then applied to all subsequent images for a given staining set, each including all four genotype/treatment conditions. Images were deconvolved (50 iterations) using the batch processing function in Huygens Essential software so that the same parameter settings were used for each immunostaining analyzed. Colocalization was quantified using Huygens Essential software which calculated Pearson’s Correlation and Manders Coefficients (M1).

### 2.12 Tissue Processing for RNA-sequencing

Immediately after completion of the electrophysiology recording, hippocampal slices were flash frozen for processing. Frozen sections were thawed in Qiazol for lysis, and total RNA was extracted using miRNeasy kit (cat # 217084) following standard protocol with on column DNase digestion (Qiagen). RNA sequencing libraries were prepared with 200 ng of total RNA using KAPA RiboErase Stranded RNA-Seq kit (cat # KK8483) following standard protocol (Roche). Sequencing was done at Stanford Functional Genomics Facility (Stanford, CA) on an Illumina HighSeq 4000 targeting ∼40 million 150 bp paired-end reads per sample.

### 2.13 RNA-sequencing Data Processing Pipeline

FASTQ files generated from 18 samples consisted of stimulated hippocampal slices in the following four experimental groups: WT-Veh-TBS, n = 4 slices; WT-BD10-2-TBS, n = 4 slices; APP-Veh-TBS, n = 4 slices; and APP-BD10-2-TBS, n = 6 slices. Paired-end FASTQ files) were run through a unified RNA-Seq processing pipeline based on the ENCODE RNA-seq pipeline [34, 35]. FASTQ files were trimmed for adapter sequences and low quality base calls (Phred < 30 at ends) using Cutadapt v1.8.1 [36]. Trimmed reads were then aligned to the GRCm38.p6 (mm10) reference genome with STAR 2.7.6a [37] using gene annotations from GENCODE release vM23. Gene counts were calculated using RSEM v1.3.1 [38]. Quality control metrics were calculated using featureCounts v1.6 [39], PicardTools v2.23.4 [40], and Samtools v1.11 [41].

### 2.14 RNA-Seq Quality Control and Normalization

RNA-Seq quality control and normalization expected counts were compiled from gene-level RSEM quantifications and imported into R [42] for downstream analyses. Expressed genes were defined as genes with TPM > 0.2 in at least 50% of samples from each group (WT-Veh-TBS; WT-BD10-2-TBS; APP-Veh-TBS or APP-BD10-2-TBS). Only expressed genes were included in the analysis. A total of 23,407 expressed genes were used in the downstream analysis using the GENCODE M23 annotation gtf file. Outliers were defined by the majority of 3 analyses: Standardized sample network connectivity |Z scores| > 2, as previously described [43] (**Fig. S2B**); Principal component analysis was calculated and plots were colored by drug treatment and genotype effect; and hierarchical clustering using Poisson distance with PoiClaClu 1.0.2.1 (**Fig. S2A**) [44]. One sample, BD10_2_30, failed both PCA and Poisson clustering and was removed as an outlier.

### 2.15 Covariate Selection

We compiled a set of 152 RNA-Seq quality control metrics from the outputs of cutadapt, STAR, featureCounts and PicardTools (MarkDuplicates, CollectGcBiasMetrics, CollectAlignmentSummaryMetrics, CollectInsertSizeMetrics, CollectRnaSeqMetrics). These measures were summarized by the top 36 principal components, which explained the total variance of each group. Multivariate Adaptive Regression Splines (MARS) implemented in the earth package in R was used to determine which covariates to include in the final differential expression model. The potential covariates included: genotype, LTP effect, drug treatment and seqPCs. These covariates were incorporated into the earth model along with gene expression data. The model was run using linear predictors and otherwise default parameters. This model fits a maximum of 1,000 genes simultaneously and we performed 1,000 permutations randomly. None of these covariates were significant and therefore were not included in the model.

### 2.16 Differential Gene Expression & Pathway Enrichment

Differential Gene Expression (DE) analyses was performed using DESeq2 [45]. DE results with padj < 0.05 were used for downstream enrichment analysis. The biomaRt v3.12 [46] package in R was used to extract gene names, gene biotypes and gene descriptions. Enrichment for Gene Ontology (GO; biological process, cellular component and molecular function), KEGG pathways and Reactome were performed using gprofiler2 v0.2.0 [47]. Background was restricted to the expressed set of genes. Only pathways containing less than 1,000 genes were assessed. An ordered query was used, ranking genes by p-value for DGE analyses.

### 2.17 Network Co-expression Analysis

Network analysis was performed with the Weighted Gene Correlation Network Analysis (WGCNA, version 1.70 [48] package using signed networks. A soft-threshold power of 10 was used to achieve approximate scale-free topology (R^2^ > 0.8). The ‘blockwiseModules’ function was used to construct the networks. The network dendrogram was created using average linkage hierarchical clustering of the topological overlap dissimilarity matrix (1-TOM). The hybrid dynamic tree-cutting method was used to define modules. Modules were summarized by their first principal component (ME, module eigengene) and modules with eigengene correlations > 0.9 were merged. Genes within each module were ranked based on their module membership (kME), defined as correlation to the module eigengene.

### 2.18 Cell Type & Human-Mouse Co-expression Module Enrichment Analysis

Cell type enrichment analyses were performed using several mouse derived cell type-specific expression datasets [49–51]. Human-mouse co-expression modules from Wan et al. [52] were accessed from the Sage Bionetworks Synapse Portal (syn10309369.1), and grouped according to the reported consensus clusters and brain regions: yellow – A – Astroglial-like modules; light blue – B – Microglial-like modules; maroon – C – Neuronal-associated modules; green – D – Oligodendroglial-like modules; dark brown – E – Accessory neuronal associated modules. Enrichment was performed for cell type specific marker genes using Fisher’s exact test, followed by FDR-correction for multiple testing. Jaccard indices and odds ratios for Fischer’s exact tests were generated using the GeneOverlap package (1.26.0) [53].

### 2.19 Additional Statistical Analyses

Statistical significance was determined with GraphPad Prism software v9.01 (San Diego, California USA). The number of samples, mice, and measurements are included in the figure legends. Data were assessed for normal distribution and parametric or non-parametric tests were applied accordingly. Specific statistical methods for each study, including ANOVA and post hoc analyses, are specified in the figure legends.

## 3 RESULTS

### 3.1 BD10-2 promotes cell survival via TrkB and TrkC

Our laboratory previously identified LM22B-10 as a selective, small molecule activator of TrkB and TrkC that increased hippocampal cell survival and prevented dendritic spine loss in aged mice, however it had limited blood brain barrier penetration when given orally [19]. To improve bioavailability, a derivative of LM22B-10, designated as PTX-BD10-2 (BD10-2), was developed that achieves higher brain concentrations than the parent compound [20] (**Fig. 1A**). The specificity and trophic activity of BD10-2 was examined by assessing effects on survival of 3T3-cells expressing one of four neurotrophin receptors (TrkA, TrkB, TrkC, or p75^NTR^) after treatment with BD10-2, LM22B-10, or the exogenous neurotrophins that bind to these receptors (NGF, BDNF, NT3). The 3T3 parental line does not express Trk or p75 receptors and, as expected, adding BDNF or BD10-2 to serum-free medium did not support their survival (data not shown). However, 3T3 cells engineered to stably express TrkB (**Fig. 1B**), TrkC (**Fig. 1C**) and TrkA (**Fig. 1D**) receptors exhibited robust survival responses to the cognate ligands BDNF, NT-3, and NGF, respectively. Cell survival was not significantly affected in BDNF-treated 3T3-p75 cells (**Fig.1E**). BD10-2 promoted survival of 3T3-TrkB and 3T3-TrkC cells in a dose-dependent manner and at a similar efficacy and maximal effect as the parent compound LM22B-10 and the neurotrophins, BDNF and NT3 (**Fig. 1B-C**), but had no effect on survival of TrkA- or p75-expressing 3T3 cells (**Fig. 1D-E**). To determine whether BD10-2 promoted survival of cultured hippocampal neurons, the number of neurons immunostained for III β-tubulin (Tuj1), a neuron-specific marker, was quantitated after exposure to BDNF, NT-3, LM22B-10, or BD10-2. The number of surviving hippocampal neurons was higher in cells treated with BDNF, NT3, BD10-2, and LM22B-10 compared to cells in culture medium alone (**Fig. 1F, G**). These results demonstrate that BD10-2 has survival-promoting activity with similar efficacy, although with lower potency, to that of BDNF and NT-3 and that these effects are specific to TrkB- and TrkC-expressing cells.

**Figure 1:**
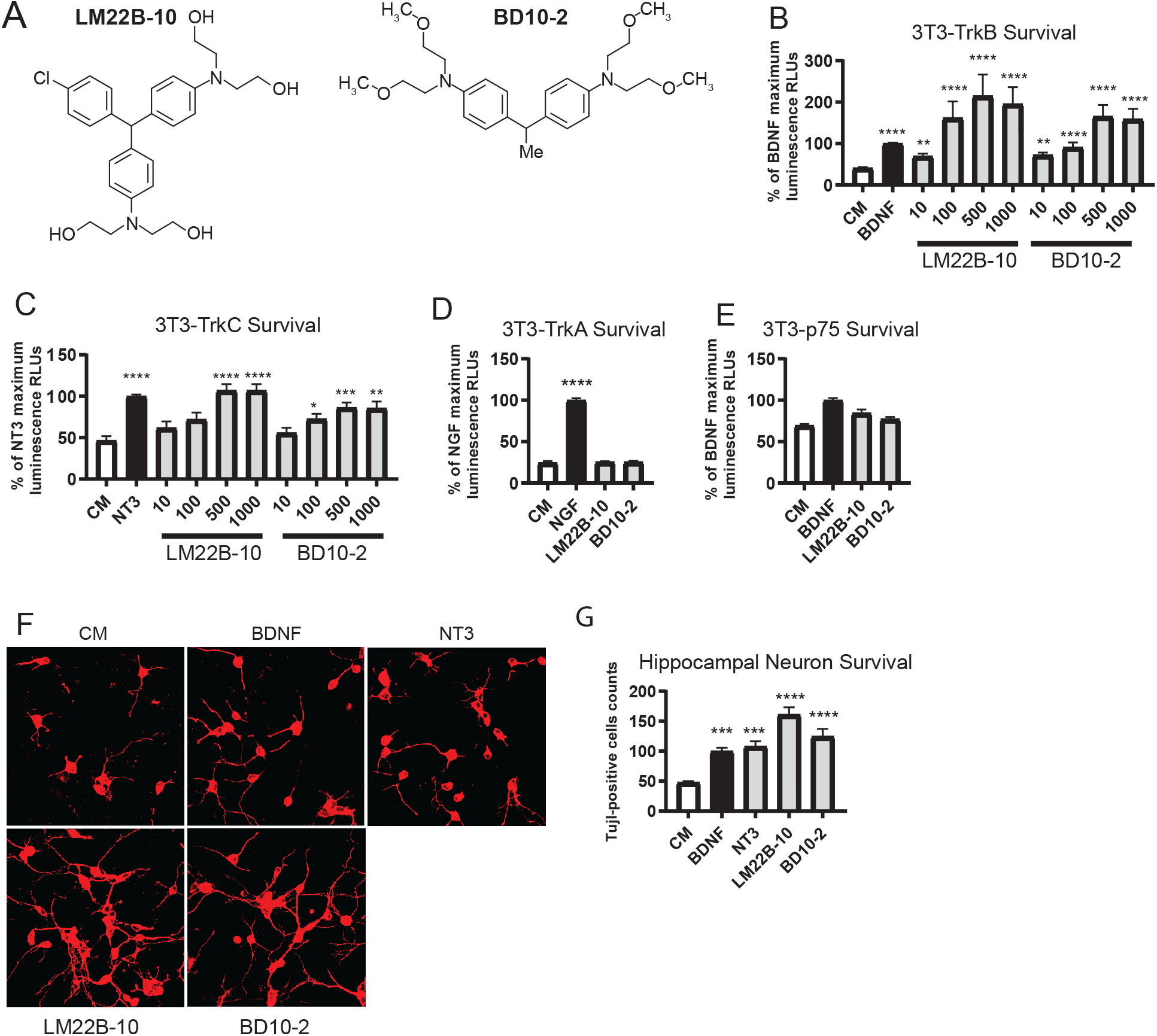
BD10-2 promotes cell survival preferentially through TrkB and TrkC. **A.** Structure of BD10-2 and its parent compound, LM22B-10. **B-E.** Trk-expressing 3T3 cells were incubated in serum-free culture media alone (CM) or in CM supplemented with exogenous neurotrophins (NTs), LM22B-10, or BD10-2 for 72-96 h until survival was measured using the ViaLight assay. BD10-2 and its parent compound LM22B-10 had a dose dependent effect on survival of **B.** 3T3-TrkB **C.** 3T3-TrkC cells (mean ± SEM, n = 16-40 wells derived from 8-9 independent experiments). Statistical significance was determined by Kruskal-Wallis test with Dunn’s post-hoc multiple comparison tests. *p ≤ 0.05, **p ≤ 0.01, ***p ≤ 0.001, ****p ≤ 0.0001 compared to CM alone. **D-E.** BD10-2 (1000 nM) and its parent compound LM22B-10 (1000 nM) had no effect on survival of **D.** 3T3-TrkA or **E.** 3T3-p75 cells compared to CM alone (mean ± SEM, n = 24 wells derived from 4 independent experiments). Statistical significance was determined by Kruskal-Wallis test with Dunn’s post-hoc multiple comparison tests. ****p ≤ 0.0001 compared to CM alone. **F.** BD10-2 supports the survival of cultured hippocampal neurons. Fluorescence photomicrographs (original magnification, ×40) of Tuj1-immunostained E16 mouse hippocampal neurons treated with either serum-free culture medium alone (CM), BDNF, NT3, LM22B-10 or BD10-2 for 48 h. **G.** Survival analysis of hippocampal neurons treated with either BDNF (20 ng/ml), LM22B-10 (1000 nM) or BD10-2 (1000 nM) for 48 h. Treated Tuj1-positive cells were counted and compared to wells receiving CM alone (mean ± SEM, n = 11 wells per group derived from 3 independent experiments). Statistical significance was determined by ANOVA with Dunnett’s post-hoc multiple comparison tests. **\*\*\***p ≤ 0.001, ****p ≤ 0.0001 compared to CM.

### 3.2 Acute and chronic BD10-2 treatment reduces synaptic plasticity deficits in hippocampal slices from APP^L/S^ mice at 16 months of age

BDNF activation of TrkB receptor signaling has a well-established role in promoting activity-dependent forms of synaptic plasticity such as hippocampal LTP [54, 55]. Recent work has shown that NT3/TrkC also modulates synaptic transmission [14]. We determined whether application of BD10-2 can acutely affect long-lasting changes in hippocampal synaptic plasticity. Hippocampal slices were prepared from APP^L/S^ and WT mice at 16 months of age and BD10-2 (100 nM) or its vehicle (Veh; HPCD) was added to the slice bath 20 min prior to theta burst stimulation (TBS). TBS was used to induce LTP because this pattern of stimulation is based on the “hippocampal theta rhythm”, a large-amplitude oscillation seen in electroencephalographic recordings in the range of 4-8Hz [56–59], and it more closely mimics in vivo physiological conditions than high-frequency stimulation (HFS) which consists of long stimulation bursts at 100 Hz [56, 60]. We used a 3xTBS protocol because, compared to a single TBS protocol, it induces a stronger and longer lasting change in synaptic transmission which allows the induction of late phases of LTP [6]. After 3xTBS, both the induction and maintenance of LTP were impaired in vehicle-treated hippocampal slices from APP^L/S^ mice (APP-Veh-TBS) compared to those from WTs (WT-Veh-TBS: 1 min 190 ± 11%, 130 min 124 ± 7%, n = 5; APP-Veh-TBS: 1 min after TBS application 152 ± 6%, 130 min 91 ±5%, n = 4; main effect of group for 2 h post-induction: *F*_1, 7_ = 11.66, *p* = 0.0112, RM-ANOVA). Bath application of BD10-2 prevented deficits in both LTP induction and maintenance in slices from APP^L/S^ mice (APP-BD10-2-TBS: 1 min 199 ± 9%, 130 min 128 ± 16, n = 5) when compared to APP^L/S^ slices exposed to Veh (*F*_1, 7_ = 7.078, *p* = 0.0325, RM-ANOVA; **Fig. 2A**). These data suggest that BD10-2 can restore deficits in hippocampal LTP in APP^L/S^ mice and may do so through acute effects of the ligand at the TrkB/TrkC receptors.

**Figure 2:**
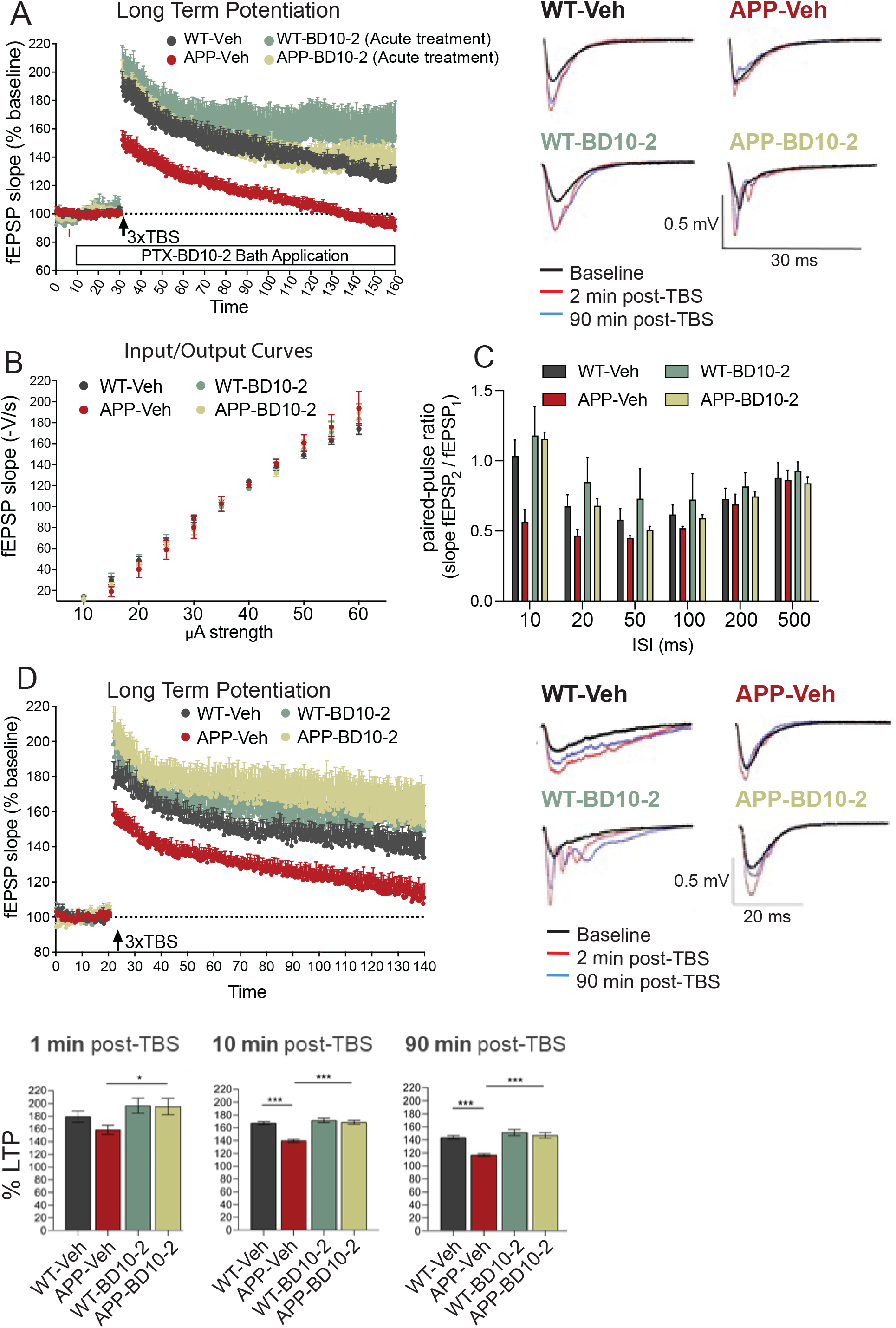
Acute and chronic BD10-2 treatment ameliorates LTP abnormalities in APP^L/S^ mice. **A.** Hippocampal slices from APP^L/S^ mice treated with vehicle (Veh) display significantly lower LTP magnitude both at the induction and maintenance phases compared to WT. Acute application of BD10-2 (100 nM) restored LTP in slices from APP^L/S^ mice. The bar along the x-axis indicates duration of drug application. Insets depict representative analog traces, obtained during baseline recording (black line), 2 min after 3xTBS (red line), and at the end of recording (130 min post 3xTBS, (blue line) for all four groups (mean ± SEM, n = 2-3 mice with 4-5 slices per condition). **B-D** Deficits in hippocampal synaptic plasticity are ameliorated in APP^L/S^ mice at 16 months of age by prior in vivo treatment with BD10-2 (mean ± SEM, n = 6-7 mice with 9-12 slices per condition). **B.** Basal synaptic transmission, as measured by input/output curves, did not differ significantly between genotypes and treatment groups. **C.** Paired-pulse ratios (PPRs) were decreased at the 10, 20 and 50 ms inter-pulse intervals in hippocampal slices from APP-Veh mice compared to WT-Veh (p = 0.0014, p = 0.0353 and p = 0.0422 respectively; one-way ANOVA with uncorrected Fisher LSD); deficits in PPRs were prevented at the 10 and 20 ms inter-pulse intervals in slices from APP-BD10-2 mice (p < 0.0001 and p = 0.0252 respectively; one-way ANOVA with uncorrected Fisher LSD). **D.** *Upper panel:* The slope of field excitatory post-synaptic potentials (fEPSPs), expressed as a % of baseline, showed a smaller initial amplitude in slices from APP-Veh mice than those from WT-Veh mice and decayed toward baseline indicating impaired maintenance of potentiation. Treating APP^L/S^ mice with BD10-2 eliminated deficits in both the induction and maintenance of LTP. *Bottom panel:* The percentage of potentiation is reduced at 10 and 90 min post-TBS in hippocampal slices from APP-Veh mice (p < 0.0001 and p < 0.0001 respectively; one-way ANOVA with uncorrected Fisher LSD) and back to WT level in slices from APP^L/S^ mice after in vivo treatment with BD10-2 at 1, 10 and 90 min post TBS (p = 0.0204, p < 0.0001 and p < 0.0001; one-way ANOVA with uncorrected Fisher LSD).

Next, we investigated whether chronic 3-month treatment with BD10-2 would ameliorate synaptic plasticity deficits in APP^L/S^ mice when the ligand was not present during recordings. Hippocampal slices were prepared from mice 24 h after they received the final dose of Veh or BD10-2, which has a plasma half-life of approximately 2 h [20]. Basal synaptic transmission, as measured by input/output curves, via stimulation of the Schaffer collateral-CA1 pathway was similar in hippocampal slices prepared from APP^L/S^ and WT mice regardless of treatment with Veh or BD10-2 (**Fig. 2B**). Pre-synaptic-mediated, short-term plasticity and synaptic vesicle release probability was investigated using paired-pulse facilitation. Paired pulse ratios (PPR) were significantly decreased in APP-Veh compared to WT-Veh mice when the time between stimulation pulses was 10 ms (p = 0.0014), 20 ms (p = 0.0353) and 50 ms (p = 0.0422; one-way ANOVA with uncorrected Fisher LSD; APP^L/S^ n = 7 mice, WT n = 5 mice; **Fig. 2C**), but not at longer inter-stimulus intervals suggesting short-term plasticity deficits. Prior BD10-2 treatment alleviated the PPR deficits in APP^L/S^ mice at 10 ms (p < 0.0001) and 20 ms (p = 0.0252; one-way ANOVA with uncorrected Fisher LSD) which did not differ from WT-Veh mice at any inter-stimulus interval suggesting that TrkB/C modulation ameliorated deficits in pre-synaptic inhibitory mechanisms. As shown in acutely prepared slices in **Fig. 2D**, LTP was also significantly impaired in hippocampal slices from chronically-treated APP-Veh-TBS mice compared to WT-Veh-TBS at 16 months of age and this finding was confirmed across 120 min of recording **(Fig. 2D)**. The potentiated responses were significantly closer to baseline in slices from APP-Veh-TBS mice compared to WT-Veh-TBS and continued to decline over the 120 min post-TBS interval (WT-Veh-TBS: 1 min 180 ± 8%, 10 min 170 ± 6%, 120 min 138 ± 10%, n = 9; APP-Veh-TBS: 1 min 158 ± 7%, 10 min 142 ± 5%, 120 min 111 ± 8%; n = 9; main genotype effect for 120 min recording post-TBS: F_1,16_ = 9.695, p = 0.0067, RM-ANOVA; **Fig. 2D**). In contrast, in slices from APP^L/S^ mice chronically treated with BD10-2, LTP was induced and maintained at a magnitude similar to WT-Veh-TBS and was significantly higher than APP-Veh-TBS mice at 1, 10 and 120-min post-TBS (APP-BD10-2: 1 min 205 ± 14%, 10 min 173 ± 12%, 120 min 151 ± 21%; n = 9; APP-Veh: 1 min 158 ± 7%, 10 min 142 ± 5%, 120 min 111 ± 8%; n = 9; main drug effect for 120 min recording post-TBS: F_1,16_ = 7.451, p = 0.0148, RM-ANOVA; **Fig. 2D**).

### 3.3 Modulation of TrkB/TrkC receptors with the small molecule ligand, BD10-2, alleviates hippocampal memory deficits in APP^L/S^ mice

Given that BD10-2, when administered either acutely to brain slices, or chronically over 3 months in vivo, prevented LTP deficits and synaptic dysfunction in hippocampal slices from APP^L/S^ mice, we determined whether chronic treatment with BD10-2 would reduce memory deficits in behavioral assays. WT and APP^L/S^ mice were treated with BD10-2 or vehicle (HPCD) by oral gavage for 3 months beginning at 13 months of age, at which time amyloid pathology, initially evident at age 3-5 months, is well-established in both the frontal cortex and hippocampus of APP^L/S^ mice [21, 22]. During the last month of BD10-2 treatment (∼15.5 months of age), hippocampal-dependent memory tests were performed including novel object recognition (NOR), novel object displacement (NOD), and Barnes maze tests to assess BD10-2’s effects on cognition. The NOR test assesses recognition memory and is based on the tendency of cognitively normal rodents to spend more time exploring a novel object than a familiar one (**Fig. 3A**). Similarly, the NOD test is based on the tendency of rodents to spend more time exploring a displaced object than one in a familiar location (**Fig. 3C**). We found that APP^L/S^ mice treated with vehicle (APP-Veh) spent significantly less time with the novel object compared to WT mice treated with vehicle (WT-Veh) (**Fig. 3B**). They also spent less time with the familiar object in a novel location in the NOD task (**Fig. 3D**). APP^L/S^ mice treated with BD10-2 (APP-BD10-2) exhibited improved ability to discriminate the novel object and location compared to APP-Veh mice (**Fig. 3B, D**). These results demonstrate that hippocampus-dependent memory deficits observed in aged APP^L/S^ mice are rescued by BD10-2.

**Figure 3:**
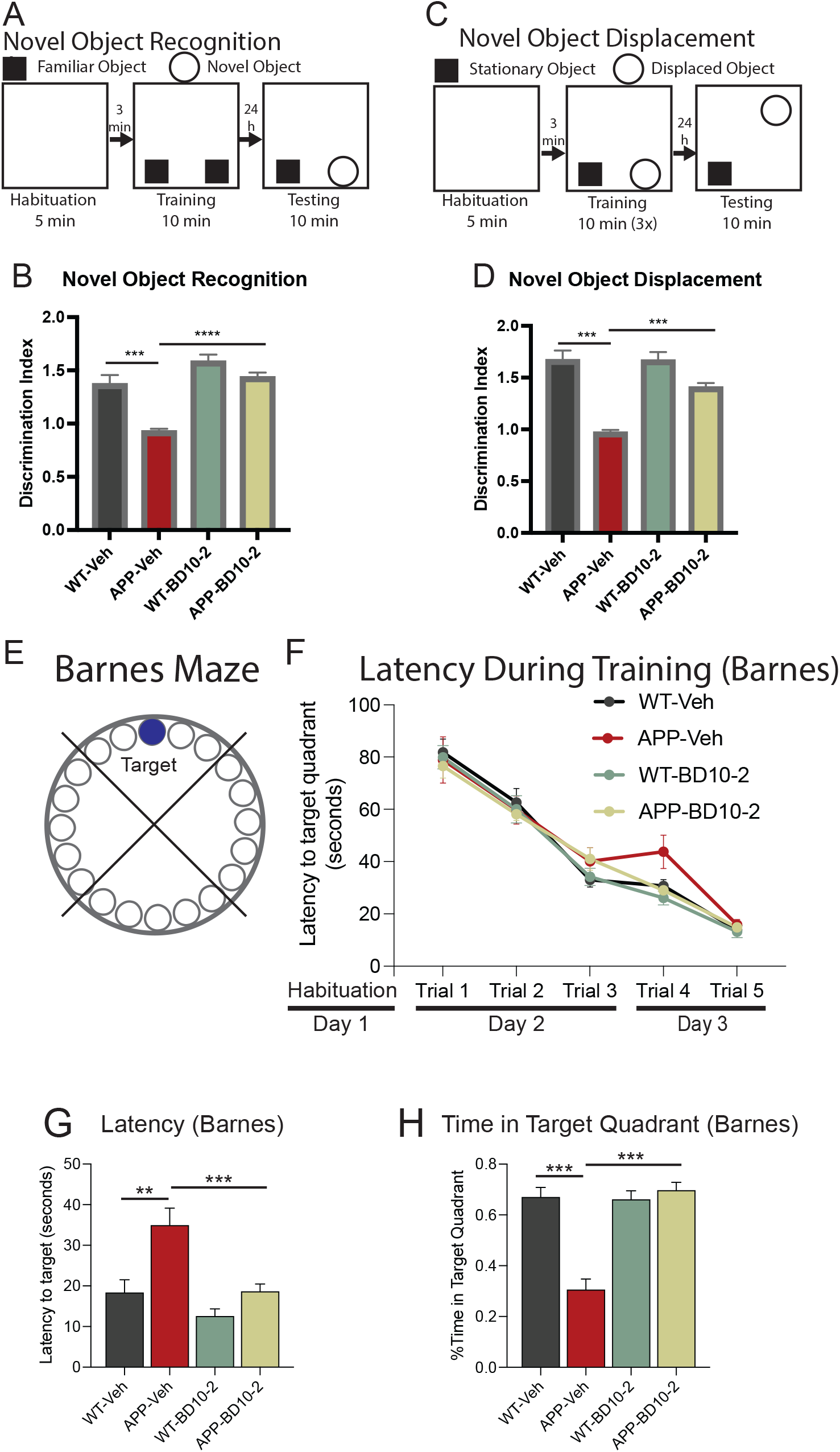
Long-term treatment with BD10-2 ameliorates cognitive abnormalities in aged APP^L/S^ mice. **A-H.** Behavioral tests of cognition in APP^L/S^ mice treated with BD10-2 by oral gavage for 3 months starting at 13 months of age (mean ± SEM, n = 9-12 mice per group). **A.** For the Novel Object Recognition (NOR) test, mice were habituated to an arena for 5 min prior to the training session where two identical objects were placed in the arena and the mice were allowed to explore for 10 min. After 24 h, mice were returned to the arena in which one of the familiar objects was replaced by a novel object and exploratory behavior around the objects was tracked for 10 min. **B.** In the NOR test, APP-Veh mice showed prominent deficits in discriminating between novel and familiar objects compared to WT-Veh (p < 0.0001; one-way ANOVA with uncorrected Fisher LSD); BD10-2 treatment prevented this deficit (p < 0.0001; one-way ANOVA with uncorrected Fisher LSD). **C.** For the Novel Object Displacement test (NOD) mice were habituated to an arena for 5 min followed by three training sessions where two identical objects were placed in the arena and the mice were allowed to explore for 10 min. After 24 h, one of the objects used during training was moved to a new location and mice were returned to the arena. Exploratory behavior around the objects was tracked for 10 minutes. **D.** In the NOD test, the ability of APP-Veh mice to discriminate a familiar object in a novel location was impaired versus WT (p < 0.0001; one-way ANOVA with uncorrected Fisher LSD); this impairment was ameliorated by BD10-2 treatment (p < 0.0001; one-way ANOVA with uncorrected Fisher LSD). **E-H.** Barnes Maze task: **E.** The Barnes maze measures spatial memory by testing a rodent’s ability to remember the location of a hole which allows them to escape mildly aversive stimuli of bright light and open space. The time spent in the target quadrant of the maze and the latency to reach the escape hole were measured. **F.** During training (Days 2 and 3), all mice learned the location of the target quadrant and escape hole as evidenced by similar latencies to the target quadrant. **G**. On the test day 4, the latency to reach the target escape hole is increased in APP-Veh mice (p = 0.0003; one-way ANOVA with uncorrected Fisher LSD), while APP-BD10-2 mice found the target significantly faster than APP-Veh mice (p = 0.0002; one-way ANOVA with uncorrected Fisher LSD). **H.** Inversely, the percentage of time spent in the target quadrant is reduced in APP-Veh mice versus WT-Veh (p < 0.0001; one-way ANOVA with uncorrected Fisher LSD); treatment of APP^L/S^ mice with BD10-2 prevented this deficit (p < 0.0001; one-way ANOVA with uncorrected Fisher LSD).

The Barnes Maze is a test of spatial learning and memory in which the mice must learn a target location using distal visual cues to escape a mildly aversive stimulus of bright light and open space (**Fig. 3E**). After habituation (day 1) and during training (days 2 and 3), all mice learned the location of the target escape hole with a similar latency (**Fig. 3F**). When tested 24 h later, APP-Veh mice took longer to enter the target escape hole than WT-Veh mice (**Fig. 3G**) and spent significantly less time in the target quadrant of the maze (**Fig. 3H**) suggesting spatial memory impairments. Treating APP^L/S^ mice with BD10-2 ameliorated the spatial memory deficits as they located the escape hole faster and spent more time in the target quadrant compared to APP-Veh mice (**Fig. 3G, H**). These results indicate that spatial memory deficits observed in aged APP^L/S^ mice are rescued by BD10-2.

### 3.4 Treating APP^L/S^ with BD10-2 normalizes synaptic markers and synaptic plasticity-related signaling proteins in hippocampal slices

To elucidate mechanisms by which BD10-2 ameliorates synaptic dysfunction in APP^L/S^ mice, we determined whether synaptic proteins that function to maintain LTP and that are known to be affected by BDNF/TrkB or NT-3/TrkC signaling are altered in APP^L/S^ hippocampal slices, and to what extent treatment with BD10-2 might affect one or more of these proteins. Hippocampal slices were collected from each group of mice that were chronically treated with Veh or BD10-2 that either received or did not receive TBS (8 treatment groups in total). Western blotting for the post-synaptic marker, PSD-95, or the pre-synaptic marker, synaptophysin, were performed using lysates prepared from hippocampal slices that did not receive TBS. APP-Veh mice at 16 months of age exhibited a 31% reduction in PSD95 levels compared to WT-Veh mice; BD10-2 prevented this decrease and showed PSD95 levels similar to WT-Veh (**Fig. 4A, B**). BD10-2 did not affect PSD-95 levels in WT mice (**Fig. 4A, B**). Although a previous study showed decreases in synaptophysin in the hippocampus of APP^L/S^ mice at 5-7 months of age [21], we observed no changes in synaptophysin (**Fig. 4A, C**).

**Figure 4:**
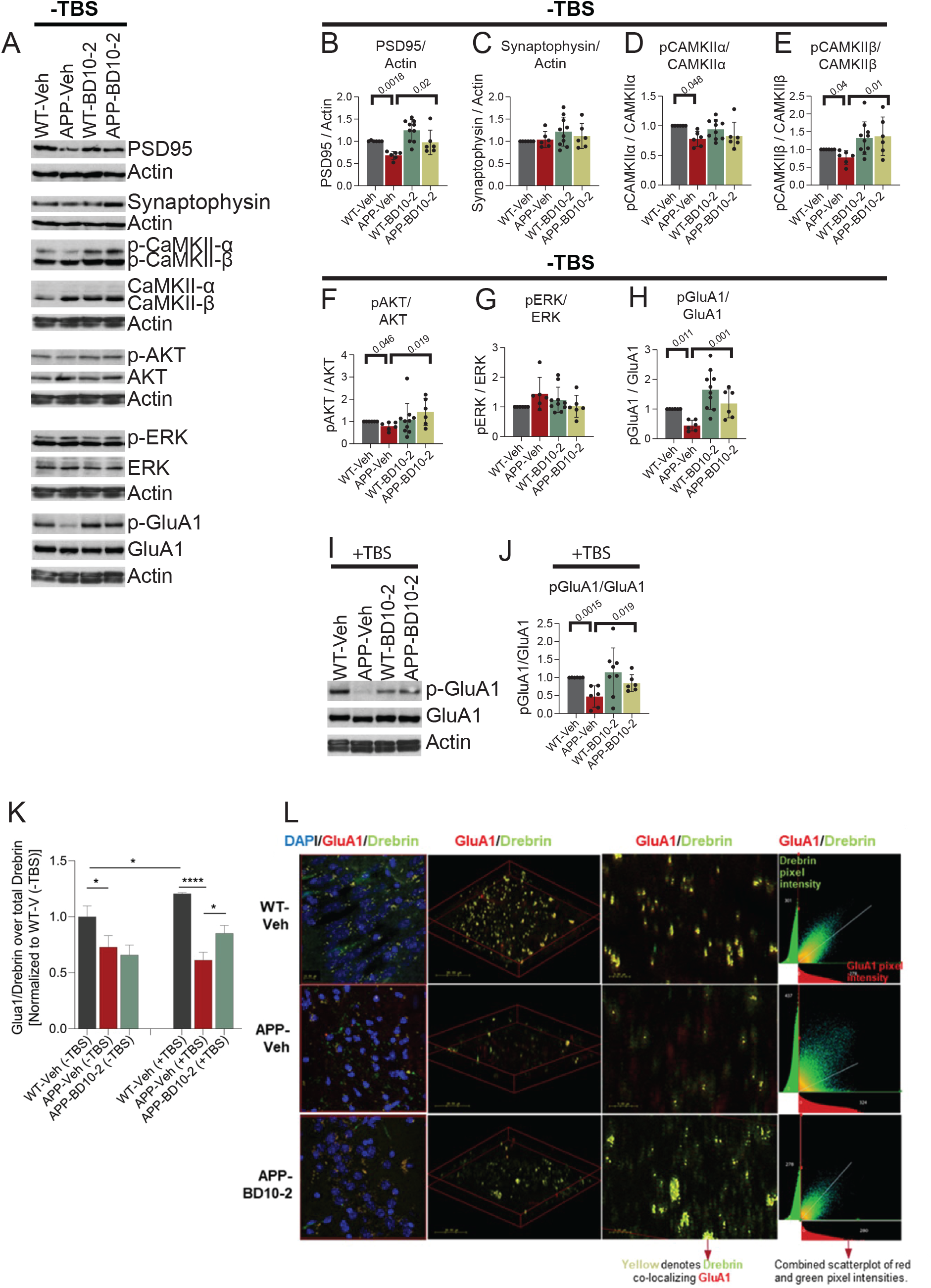
BD10-2 enhances baseline CAMKII, GluA1, and AKT phosphorylation as well as co-localization of GluA1 and drebrin in slices that underwent LTP. BD10-2 or vehicle was administered by oral gavage to 13-month-old APP^L/S^ mice for three months, followed by collection of hippocampal slices. **A.** Representative western blots of unstimulated (TBS-) hippocampal slices. **B-H, J.** Western blots of hippocampal slice extracts were quantitated by determining the ratios of phospho (p) protein over total protein or total protein over actin, and then normalized to respective WT-Veh groups. Statistical significance was determined using either one way ANOVA with Sidak post hoc test or Kruskal-Wallis test with post-hoc Dunn’s multiple comparisons test (mean ± SEM, Sample size = 6-10 hippocampal slices, 3-5 mice per group, with two independent western blots averaged per slice). **B-H** Quantification of unstimulated (TBS-) hippocampal slices shown in A. **I.** Representative western blot for hippocampal slices that underwent 3xTBS. **J.** Quantification of stimulated (TBS+) hippocampal slices shown in I. **K-L.** Immunofluorescence staining analysis of co-localization between GluA1 and drebrin in slices that underwent 3xTBS. **K.** Mander’s Coefficient (M1) for samples stained with GluA1 and drebrin. All samples are normalized to WT-Veh without TBS. Proportion of GluA1 colocalized with drebrin in APP and WT mouse hippocampal slices without and with stimulation (mean ± SEM, n = 3 mice per group). *p < 0.05; ***p < 0.0005; ****p < 0.0001 by Uncorrected Fisher’s LSD multiple comparison following one-way ANOVA. **L.** Representative 63x confocal microscope z-stack images of stained DAPI cells and colocalized drebin and GluA1 (yellow) from different angles in slices that underwent 3xTBS. The CA1 region of the hippocampus was photographed with 0.3 μm step size to create a composite image of each slice. Middle panel demonstrates decreased colocalization in the APP vehicle mouse as compared to the WT vehicle and the transgenic drug condition. Lower middle panel demonstrates increased colocalization for the APP^L/S^ mouse treated for 3 months with BD10-2 as compared to the untreated APP^L/S^ mouse.

LTP induction is mediated, in part, by AMPA receptor (AMPAR) trafficking in postsynaptic neurons [61]. AMPAR delivery to the synapse can be initiated by phosphorylation of the AMPAR GluA1 subunit by CAMKII [62]. BDNF/TrkB signaling has been implicated in this process through activation of PLCγ/PKC signaling which activates CAMKII [62–66]. We assessed via western blotting the effects of BD10-2 on the phosphorylation and activation of kinases associated with three of the main TrkB/C signaling pathways including: CaMKII α/β (phosphorylated (p) at site Thr286), a signaling intermediate of the PLCγ/PKC pathway; AKT (Ser 473) part of the PI3K/AKT pathway; and ERK1/2 (Thr202/Tyr204 site) in the MAPK/ERK pathway. Activation of each of these pathways has been shown to be disrupted in AD mice contributing to LTP deficits [67–73]. First, we assessed whether activation, as indicated by phosphorylation of selected signaling proteins within these pathways, was affected in APP^L/S^ mice under baseline conditions (TBS-) and, if so, whether BD10-2 normalizes these changes. Activity levels, as indicated by the ratio of pCaMKIIα, pCaMKIIβ and pAKT over respective total protein were significantly decreased in slices from APP-Veh mice compared to WT-Veh slices (**Fig. 4A, D-F**), while the ratio of pERK was unchanged (**Fig. 4A, G**). Chronic treatment with BD10-2 normalized pCaMKIIβ and pAKT ratios in APP hippocampal slices (**Fig. 4A, E, F**), but not pCaMKIIα (**Fig. 4A, D**) or pERK ratios (**Fig. 4A, G**). Phosphorylation of CaMKIIα, CaMKIIβ, AKT and ERK were unchanged in WT-BD10-2 mice compared to that from WT-Veh mice (**Fig. 4A, D-G**). Total protein levels, as indicated by ratio of protein over actin, of each of these signaling molecules did not differ between any of the treatment groups (**Fig. 4A, S1A-D**). Since CAMKII can phosphorylate GluA1 and thereby regulate its delivery to the synapse [66, 74], we assessed the effects of BD10-2 on GluA1 phosphorylation at Ser831 (pGluA1), a site phosphorylated by CaMKII [75]. The ratio of pGluA1/GluA was reduced in hippocampal slices from APP-Veh mice compared to WT-Veh while BD10-2 prevented this decrease. The ratio pGluA1/GluA1 did not change in WT mice in response to BD10-2 treatment (**Fig. 4A, H**) and total protein levels of GluA1 remained stable in all treatment conditions (**Fig. 4A, S1E**).

Since we observed changes in baseline activation of proteins associated with LTP that were normalized by BD10-2, we asked whether the same would be true in hippocampal slices that underwent TBS (TBS+). Like unstimulated hippocampi, TBS+ slices from APP-Veh mice showed reductions in PSD-95 and pAKT/AKT compared to WT-Veh mice, however BD10-2 had no effect on these deficits (**Fig. S1F, G, M**). Notably, pGluA1/GluA1 was also reduced in TBS+ hippocampal slices from APP^L/S^ mice compared to WT-Veh; BD10-2 prevented this decrease (**Fig. 4I, J**). No changes in any of the other activation ratios or protein levels examined in the unstimulated slices were seen between any of the treatment groups in TBS+ slices (**Fig. S1F, H-L, N-Q**).

Since BD10-2 normalized pGluA1 levels in both stimulated and unstimulated hippocampal slices of APP^L/S^ mice and GluA1 phosphorylation is associated with AMPAR transport to the synapse, we assessed whether chronic BD10-2 treatment increased levels of GluA1 in synapses within the CA1 region of hippocampal slices from APP^L/S^ and WT mice with and without TBS. Synaptic localization was evaluated employing the colocalization of immunostaining for GluA1 and the postsynaptic marker, drebrin. To compare the effects of stimulation, genotype, and BD10-2, data was normalized to the WT-Veh condition without TBS. At baseline conditions (without TBS), the amount of GluA1 colocalized with drebrin (M1 Mander’s coefficient) was significantly decreased in hippocampal slices from APP-Veh compared to WT-Veh mice; BD10-2 did not change colocalization in unstimulated slices (**Fig. 4K, L**). GluA1 colocalization with drebrin was increased in WT-Veh slices 2 h post-TBS compared to unstimulated slices from WT-Veh mice. In contrast, TBS failed to increase GluA1-drebrin colocalization in APP-Veh slices compared to unstimulated APP-Veh slices (**Fig. 4K, L**). Similar to the unstimulated condition, GluA1-drebrin colocalization was reduced in slices from stimulated APP-Veh slices compared to stimulated WT-Veh slices. Interestingly, in contrast to unstimulated slices, BD10-2 partially rescued GluA1-drebrin colocalization in stimulated hippocampal slices from APP^L/S^ mice (**Fig. 4K, L**), suggesting that one mechanism by which BD10-2 might alleviate LTP deficits is by enhancing localization of GluA1-AMPARs to synapses.

### 3.5 Altered activity-dependent transcriptional signatures observed in APP^L/S^ mice are normalized by BD10-2 treatment

Synaptic activity-dependent transcription is critical for stabilizing the long-lasting changes established in late phase LTP (L-LTP), which requires the synthesis of new proteins implying a transcriptional shift at some point during potentiation [5, 76–84]. We examined whether TBS-driven LTP affects gene transcription in WT and APP^L/S^ mice and if the restorative effects of BD10-2 on LTP in APP^L/S^ mice were associated with L-LTP related transcriptional changes. To this end, RNA sequencing was performed on hippocampal slices from WT and APP^L/S^ mice that were chronically treated with BD10-2 or vehicle and used for the LTP experiments in **Fig. 2D**. A principal component analysis of the whole transcriptome revealed a strong genotype and TBS effect as there was a clear separation between their respective principal component groupings (**Fig. 5A**) and volcano plots (**Fig. S3**). To identify L-LTP-associated gene expression, we examined the effects of TBS on differential gene expression (padj < 0.05) for groups that exhibited LTP (WT-Veh-TBS vs WT-Veh; APP-BD10-2-TBS vs APP-BD10-2) or impaired LTP (APP-Veh-TBS vs APP-Veh) (**Fig. 5B-D, Tables S1-S7**). Genes that changed with TBS in WT-Veh mice, which exhibited LTP, and APP-Veh mice, which did not, can be classified by quadrant. The y-axis shows the difference in TBS gene expression between WT-Veh and APP-Veh mice, with opposing direction of fold change indicating those associated with LTP (**Fig. 5B**). Quadrants 1 and 4 show genes that are upregulated with TBS in WT mice. Both show strong general immune-associated Gene Ontology (GO) enrichment (**Table S7**). Quadrant 4 gene enrichments represent those lessened or lost in the APP-Veh-TBS mice, these include cytokines, and genes involved in apoptosis and MAPK signaling. Genes in quadrant 2 are downregulated with TBS in WT-Veh mice and are increased or upregulated with TBS in APP-Veh mice. Quadrant 2 genes show relevance to synaptic function, signaling, angiogenesis, and neuron projection morphogenesis. Taken together, these data suggest that activity-dependent LTP associated transcriptional changes involve synaptic and immune processes. Additionally, the similarity in transcription patterns between the two groups that exhibited activity dependent LTP after TBS (WT-Veh, and APP-BD10-2) (**Fig. 5**), supports the hypothesis that BD10-2 reduces alterations in activity-dependent LTP associated gene expression in APP^L/S^ mice.

**Figure 5:**
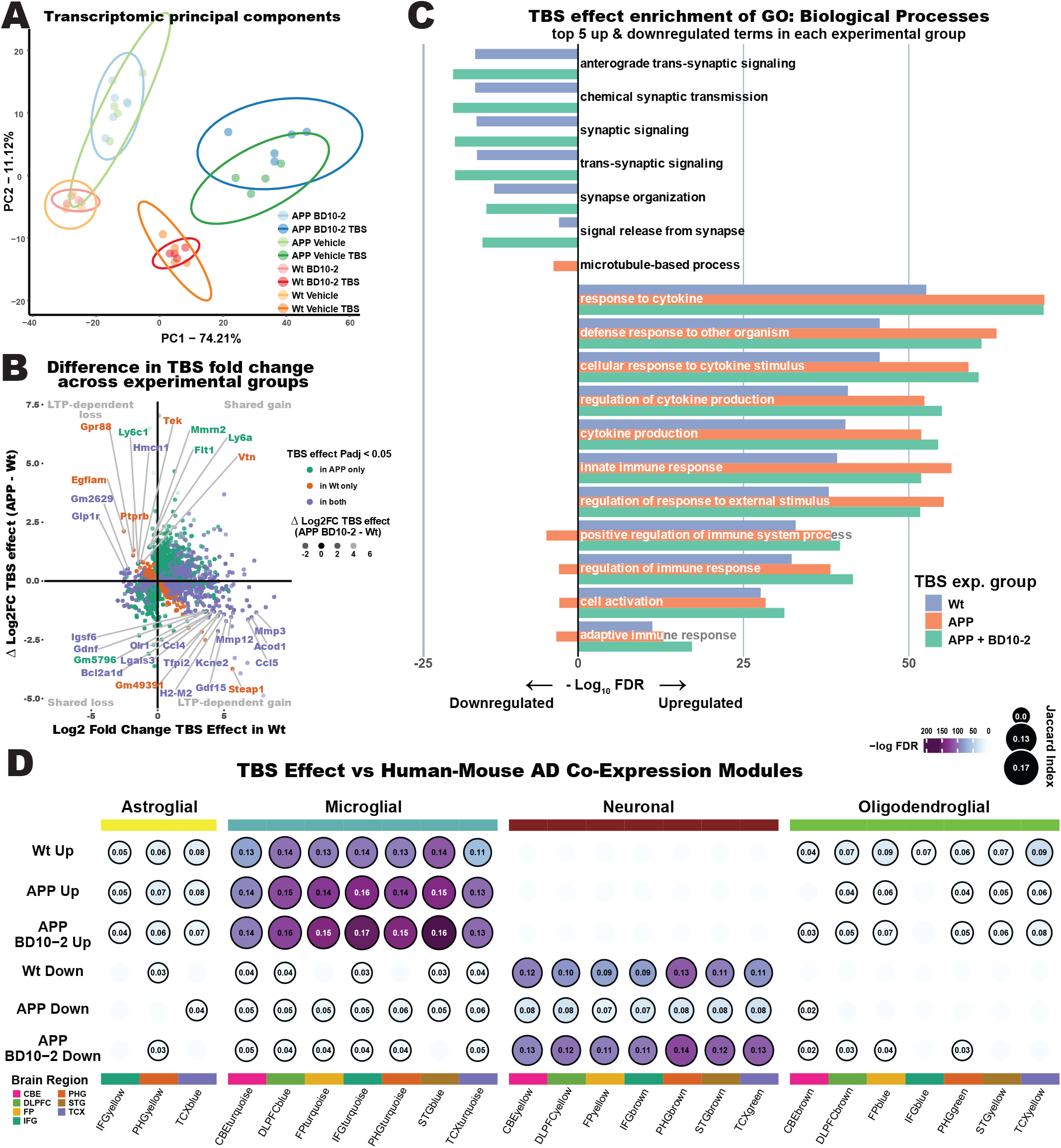
Theta burst stimulation (TBS) causes activity-dependent transcriptional changes in WT mice that are absent in APP^L/S^ mice and rescued by BD10-2 treatment. **A.** Principal component analysis of the top 1000 most variable genes in hippocampal slices from mice in each of the eight treatment groups: WT-Veh (Orange), WT-BD10-2 (Red), APP-Veh (Green), APP-BD10-2 (Blue) either with TBS (darker shade) or sham TBS (lighter shade). **B.** DE fold change comparison between TBS effects in the three treatment groups: WT-Veh (WT-Veh-TBS vs WT-Veh), APP-Veh (APP-Veh-TBS vs APP-Veh), and APP-BD10-2 (APP-BD10-2-TBS vs APP-BD10-2). Genes shown are limited to those significantly DE (padj < 0.05) in both WT-Veh and APP-BD10-2 together (those displaying normal LTP) or APP-Veh (which failed to induce and maintain LTP). The x-axis depicts TBS effect in WT-Veh group, while the y-axis shows the difference in log_2_ fold change between APP-Veh and WT-Veh groups, indicating LTP-relevant discordance in TBS expression. Genes significant (padj < 0.05) in respective groups (or their overlap) are color coded. Spot transparency is graded by the difference in log_2_ fold change between WT-Veh and APP-BD10-2 groups, indicating discordance in TBS expression between the two groups that display intact LTP. **C**. Gene Ontology enrichment showing the effect of TBS in the three treatment groups. Up or downregulated genes (padj < 0.05) for TBS in each group was tested for enrichment of GO: Biological Process, and the top 5 terms for each directional set in each group were merged into 18 total terms. **D.** Overlap enrichment (Fisher’s exact) of AD-related human-mouse co-expression modules from Wan et al. [52] compared with upregulated (top 3 rows)/downregulated (bottom 3 rows) genes in the TBS effect in the three treatment groups (padj < 0.05). Circles are sized and annotated with the Jaccard index of their overlap and are colored by the significance of the overlap enrichment (-log FDR; circles with an FDR < 0.05 have a bold edge). Modules are grouped based on “consensus clusters”, yellow, Astroglial-like modules; light blue, Microglial-like modules; maroon, Neuronal-associated modules; green, Oligodendroglial-like modules. Modules are additionally annotated along the bottom by the associated brain region of the human AD cohort from which they were derived, CBE, cerebellum; DLPFC, dorsolateral prefrontal cortex; FP, frontal pole; IFG, inferior frontal gyrus; PHG, parahippocampal gyrus; STG, superior temporal gyrus; TCX, temporal cortex.

To further explore these LTP-associated biological processes, we examined the top GO processes associated with genes that were upregulated or downregulated with TBS across the three experimental groups: WT-Veh, APP-Veh, APP-BD10-2 (**Fig. 5C, Tables S4-S6**). Looking at the top-enriched processes in downregulated genes, GO pathways associated with synapse-related processes such as ‘synaptic signaling’, ‘synapse organization’, and ‘signal release from synapse’ show a notable activity-dependent pattern. This TBS-associated downregulation only occurred in the groups demonstrating LTP (WT-Veh-TBS and APP-BD10-2-TBS) and was absent in APP-Veh slices which exhibited impaired LTP (**Fig. 5C**). Examining the top GO processes enriched in genes upregulated by TBS showed that immune system processes were represented across all groups. Many immune-related genes were upregulated with TBS in WT-Veh mice; however, the magnitude of enrichment was greater in both APP-Veh and APP-BD10-2 groups. This result may reflect an upregulated immune response to amyloid pathology in APP-Veh that is not addressed by BD10-2 treatment.

The relevance of the effect of TBS on gene expression in AD mice to human AD was examined by performing an overlap enrichment analysis between DE genes associated with TBS in our mouse study (padj < 0.05) and an AD-related human-mouse co-expression module study by Wan et al. [52]. Wan’s group examined consensus modules of genes that are altered in brains of human AD patients (versus non-AD controls) across multiple studies ROSMAP [85], MSSM [86], and Mayo [87] and brain regions gathered by the Accelerating Medicines Partnership-Alzheimer’s Disease (AMP-AD) Consortium. These co-expression modules share sizeable overlap with a broad range of mouse models of AD (251 DE gene sets), providing strong evidence they contain cross-species AD-associated genes. The modules largely group into consensus clusters that correlate to cell types. By comparing these clusters with our DE gene sets (padj < 0.05), we determined which AD-related modules might be enriched in genes that are differentially expressed after TBS. The analysis demonstrated significant overlap enrichment of AD-relevant neuronal modules with genes that were downregulated by TBS in both WT and APP-BD10-2 mice but showed notably reduced enrichment in both overlap size and significance in the APP group (**Fig. 5D, Fig. S4A**), consistent with activity-dependent LTP associated enrichment. In contrast, microglial consensus clusters, which correspond to immune associated processes, showed consistent enrichment of genes upregulated by TBS across all treatment groups with slightly higher enrichment in both APP and APP-BD10-2, similar to **Fig. 5C**. This suggests that upregulation of this microglial module is relevant to amyloid model mice and human AD but occurs regardless of whether or not samples exhibited activity-dependent LTP [88]. Oligodendroglial and astroglial modules demonstrated a low level of enrichment after TBS and showed little evidence for module level activity-dependent enrichment. Noting that L-LTP transcriptional effects are measured 130 minutes (during late-rather than early-stage LTP) after the initiation of TBS, the data presented in **Fig. 5** highlight that TBS alters transcription of immune, microglial, and synaptic/neuronal processes. Perhaps unexpectedly, downregulation of synaptic/neuronal processes—rather than upregulation—is consistently specific to the LTP groups, demonstrating the strongest transcriptional signature of activity-dependent LTP, which is rescued in APP^L/S^ mice with the addition of BD10-2. While broad categories of immune/microglial processes did not show clear association with activity-dependent LTP, some immune-associated genes did, suggesting more nuanced modulation of immune-associated gene expression by BD10-2 may play a role in LTP rescue.

### 3.6 BD10-2 affects genes related to immune responses, synaptic transport, microglia and oligodendrocyte function in stimulated hippocampal slices from APP^L/S^ mice

After examining DE effects caused by TBS in the experimental groups, we asked whether any DE effects are driven by APP^L/S^ genotype and whether BD10-2 can ameliorate these effects among stimulated samples. Our goal was to identify changes that might influence BD10-2 mediated rescue of LTP deficits in APP^L/S^ mice described above. To answer this, we asked three questions that can be answered by comparing three DE effect groups. First, how does gene expression change in APP^L/S^ mice compared to WT mice (APP effect: APP-Veh-TBS vs WT-Veh-TBS mice)? Second, how does gene expression change in BD10-2 treated APP^L/S^ mice compared to untreated APP^L/S^ mice (BD10-2 effect: APP-BD10-2-TBS vs APP-Veh-TBS mice)? And third, does BD10-2 treatment restore gene expression in APP^L/S^ mice to a more WT-like state (APP-BD10-2 effect: APP-BD10-2-TBS vs WT-Veh-TBS mice)? It is important to note that the APP and APP-BD10-2 effects are normalized to the same baseline group (WT-Veh-TBS) allowing relative comparison of the effects of APP genotype alone compared to the restorative effects of BD10-2 on APP^L/S^ mice in both overlap expression analysis and gene network module analysis. If BD10-2 restores APP^L/S^ transcription to resemble WT-Veh transcription, there should be relatively few DE genes in the APP-BD10-2 effect. We can compare the APP effect with the APP-BD10-2 effect to observe pathway alterations between APP-Veh-TBS mice, which do not exhibit LTP, and APP-BD10-2-TBS mice which do exhibit LTP.

In terms of principal components (**Fig. 5A**), shifts related to BD10-2 were much smaller than those due to APP genotype or TBS; unsurprising given the last dose was administered over 24 hours prior to brain harvesting. The largest BD10-2 mediated DE effect observed was in the APP-TBS mice (BD10-2 effect): [APP-BD10-2-TBS (dark blue) vs APP-Veh-TBS (dark green)] (**Fig. 5A**). However, this BD10-2 effect was smaller than either group’s within-group variance, limiting our statistical power to measure these changes confidently. Accordingly, only 8 genes exhibited a BD10-2 DE effect after multiple testing correction compared to 1,384 significant DE genes associated with the APP effect and 2,858 DE genes in the APP-BD10-2 effect after multiple testing correction (padj < 0.05) (**Fig. 5A, 6A-B, Fig. S3, Tables S8-10**). The top significant DE genes in the BD10-2 effect—including some that fall below the padj < 0.05 threshold—include several complement pathway specific genes (C3, C1qa), an actin polymerization regulator (Kank1), a pair of Annexins (Anxa2 and Anxa8), and genes associated with motility and neurite growth including Hydin and axonemal dynein (Dnah11) [89] (**Fig. 6A, B**).

**Figure 6.**
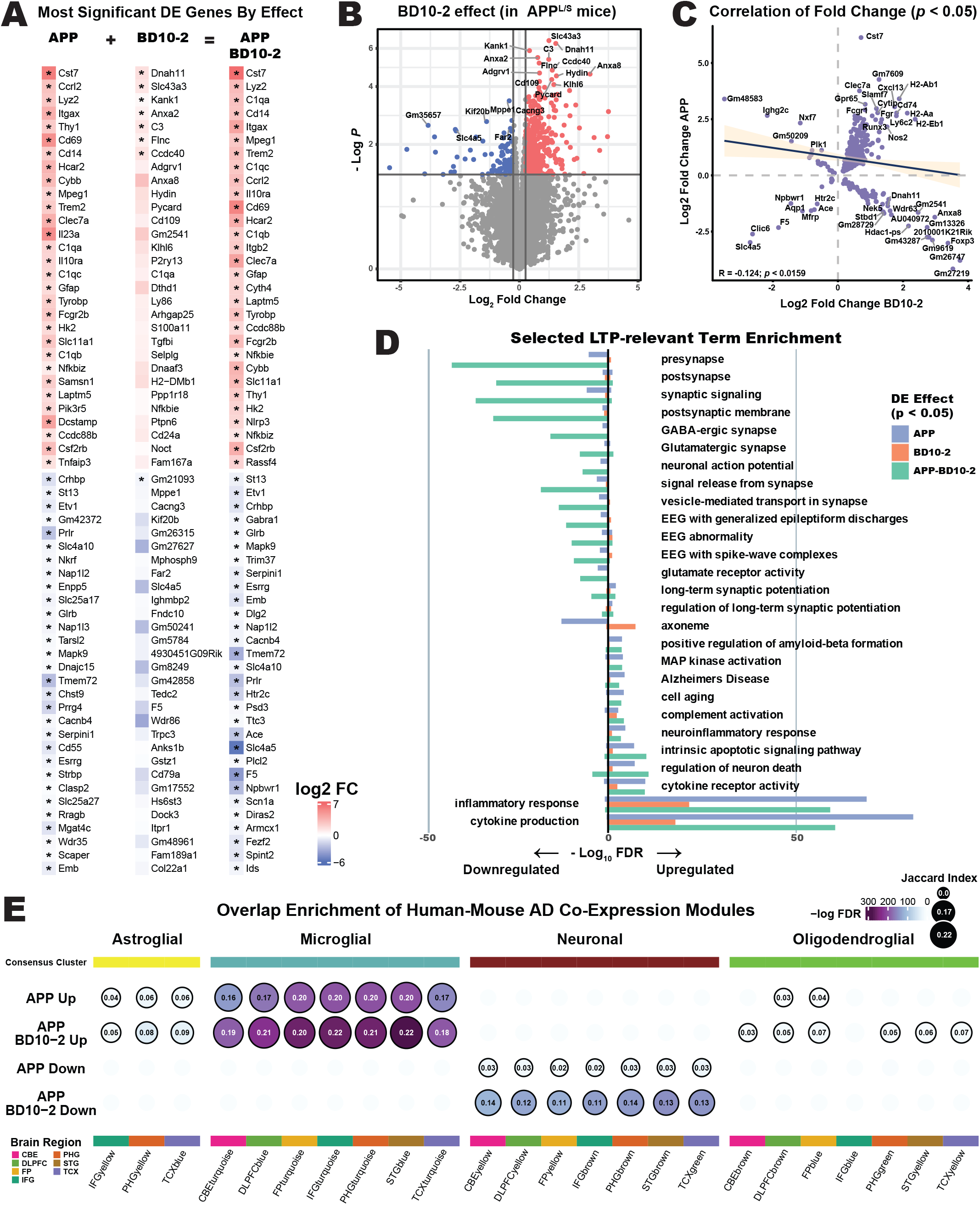
In vivo treatment of 16-month-old APP^L/S^ and WT mice with BD10-2 causes transcriptomic changes in post-TBS hippocampal slices. **A.** The top 30 upregulated and downregulated DE genes ranked by FDR adjusted p-value in each of three DE comparisons: the APP effect (APP-Veh-TBS vs WT-Veh-TBS), the BD10-2 effect (APP-BD10-2-TBS vs APP-Veh-TBS), and the APP-BD10-2 effect (APP-BD10-2-TBS vs. WT-Veh-TBS). Colored by their log_2_ fold change expression, genes with an asterisk have padj < 0.05 significance. **B.** Volcano plot showing differentially expressed (DE) genes for the BD10-2 effect (APP-BD10-2-TBS vs. APP-Veh-TBS). **C.** A correlation of fold change between the BD10-2 effect on the x-axis and the APP effect on the y-axis in genes that are nominally significant (p < 0.05) for both effects. **D.** Enrichment (g:SCS adjusted FDR) of selected ontologies/pathways relevant to LTP in downregulated (left) and upregulated (right) nominally significant (p < 0.05) DE genes for the three effect comparisons. **E.** Overlap enrichment (Fisher’s exact) of AD-related human-mouse co-expression modules from Wan et al. [52] compared with upregulated (top 2 rows)/downregulated (bottom 2 rows) DE genes (padj < 0.05) in the APP effect and the APP-BD10-2 effect. Circles are sized and annotated with the Jaccard index of their overlap and are colored by the significance of the overlap enrichment (-log FDR; circles with an FDR < 0.05 have a bold edge). Modules are grouped based on “consensus clusters”, yellow, Astroglial-like modules; light blue, Microglial-like modules; maroon, Neuronal-associated modules; green, Oligodendroglial-like modules. Modules are additionally annotated along the bottom by the associated brain region of the human AD cohort from which they were derived, CBE, cerebellum; DLPFC, dorsolateral prefrontal cortex; FP, frontal pole; IFG, inferior frontal gyrus; PHG, parahippocampal gyrus; STG, superior temporal gyrus; TCX, temporal cortex.

Many of the top upregulated genes in the APP effect were further upregulated in the APP-BD10-2 effect, and in a few cases (e.g., C1qa) appeared upregulated in the BD10-2 effect as well (**Fig. 6A**). Setting a relaxed significance threshold (p-value < 0.05), we compared the fold change of DE for the APP effect (y-axis) and BD10-2 effect (x-axis) to gauge the correlation of genes nominally significant in both (379 genes) (**Fig. 6C**). This revealed many genes in quadrant 1 which are upregulated by both effects. Quadrant 1 is enriched for inflammatory response and cytokine production (**Table S14**). Quadrant 4 contained genes that show inverse correlation of APP and BD10-2 effects: downregulated in APP^L/S^ mice and upregulated with BD10-2 treatment. Quadrant 4 was enriched for microtubule-based processes axoneme, and cilium processes. Quadrant 3 contained genes that are downregulated in both APP and BD10-2 effects and had few genes but some significant enrichment for anion transport/signaling. Although few genes were shared between APP and BD10-2 effects, we observed a notable concordance of upregulated inflammation and downregulated synaptic function-related signaling pathways.

Next, we examined selected LTP-relevant pathways including synaptic function, axonemal, and immune terms informed by our analysis of DE effects caused by TBS. A standard pathway enrichment analysis was performed on APP, BD10-2, and APP-BD10-2 effects (**Fig. 6D, Tables S11-S13**). A relaxed p-value threshold of nominally significant genes (p < 0.05) for all groups in this comparison was applied to be able to observe BD10-2 effect changes for comparison. Across several synaptic, signaling, and potentiation related terms, the APP-BD10-2 effect showed a much larger enrichment in downregulated genes than APP alone (**Fig. 6D, Tables S11-S13**), consistent with the activity-dependent LTP associated downregulation seen in **Figure 5**. Upregulated genes in both APP and APP-BD10-2 effects were enriched in pathways associated with inflammatory response, cytokines, apoptosis, and MAPK activation. The APP-BD10-2 effect showed a mild decreased enrichment in inflammatory response and cytokine production, and a mild increased enrichment of complement and apoptotic processes, compared to the APP effect (**Fig. 6D**). For most pathways, enrichment associated with the BD10-2 effect was small and often in the same direction as the APP and APP-BD10-2 effects. Notably the axoneme pathway, which contains several motility and cytoskeletal genes, was one of the few pathways where enrichment associated with the APP effect was directly opposed by a pronounced BD10-2 effect, and where no enrichment was observed in APP-BD10-2 agreeing with the correlation in individual genes in quadrant 4 of **Figure 6C** (**Fig. 6D, Tables S11-S13**).

An overlap enrichment analysis was performed to determine if DE genes (padj < 0.05) associated with the APP effect and the APP-BD10-2 effect were enriched for AD-relevant human-mouse co-expression modules [52]. This analysis revealed that neuronal human-mouse AD co-expression modules showed a mild enrichment in APP effect downregulated genes and a distinctly greater enrichment in APP-BD10-2 effect downregulated genes (**Fig. 6E, Fig. S4B**). This suggested that transcriptional changes observed in APP and APP-BD10-2 mice compared to WT-Veh mice occur in AD relevant genes. Microglial AD-relevant modules were strongly enriched in genes upregulated in both the APP effect and the APP-BD10-2 effect (**Fig. 6E**). This finding demonstrates that BD10-2 and genotype both upregulate a number of broad scale immune processes regardless of whether the sample exhibited activity-dependent LTP. Upregulated genes in astroglial and two oligodendroglial modules (prefrontal cortex and prefrontal pole) showed debatably higher enrichment in the APP-BD10-2 effect than in the APP effect alone, although four additional oligodendroglial modules showed significant enrichment only in APP-BD10-2 effect upregulated genes, suggesting BD10-2 may be activating oligodendroglial processes (**Fig. 6E**). Together, we found that long-term dosing with BD10-2 imparts transcriptomic shifts in neuronal, microglial, and oligodendroglial, human-mouse AD co-expression modules that are otherwise smaller or absent when examining APP effect alone.

Co-expression modules in the DE gene sets were independently derived in order to examine their relationship to AD pathology using a weighted gene co-expression network analysis (**Fig. 7**). This module-based assessment offers an additional approach for detection of shared expression changes associated with BD10-2 compared to assessment of the enrichment of individual DE genes. There were 23,407 genes expressed in hippocampal slices from the four groups: WT-Veh-TBS, APP-Veh-TBS, WT-BD10-2-TBS and APP-BD10-2-TBS. Among these, 34 modules were detected and 28 of these had a minimum size of 50 genes (**Fig. 7A,B**). Using the same effect comparisons that were used for the overlap enrichment analysis in **Fig. 6E** (the APP effect and the APP-BD10-2 effect), there were 5 modules significantly associated with the APP effect and 8 modules associated with the APP-BD10-2 effect (FDR < 0.05). Some of the strongest AD-related modules were significantly upregulated in both the APP effect and the APP-BD10-2 effect (red in upper panel of **Fig. 7B, Fig. S2A**). These modules showed more upregulation for the APP-BD10-2 effect versus the APP effect alone, echoing the above analyses. The three modules with the strongest upregulation (M2, M6, M18) had cell type-specific enrichment for microglia and astrocytes using an independent mouse cortex and hippocampus single-cell data set from Zhang et al [51] (green in lower panel of **Fig. 7B**). Module M6 showed a particularly strong effect and contained many of the top DE genes from our analysis of individual gene expression changes (**Fig. 6A**), including C1q subunits, Trem2, Gfap, and Tyrobp (**Fig. 7C**). Looking at pathway/ontology enrichment for highlighted modules in **Fig. 7D**, we see M6 enriched for immune-associated function.

**Figure 7.**
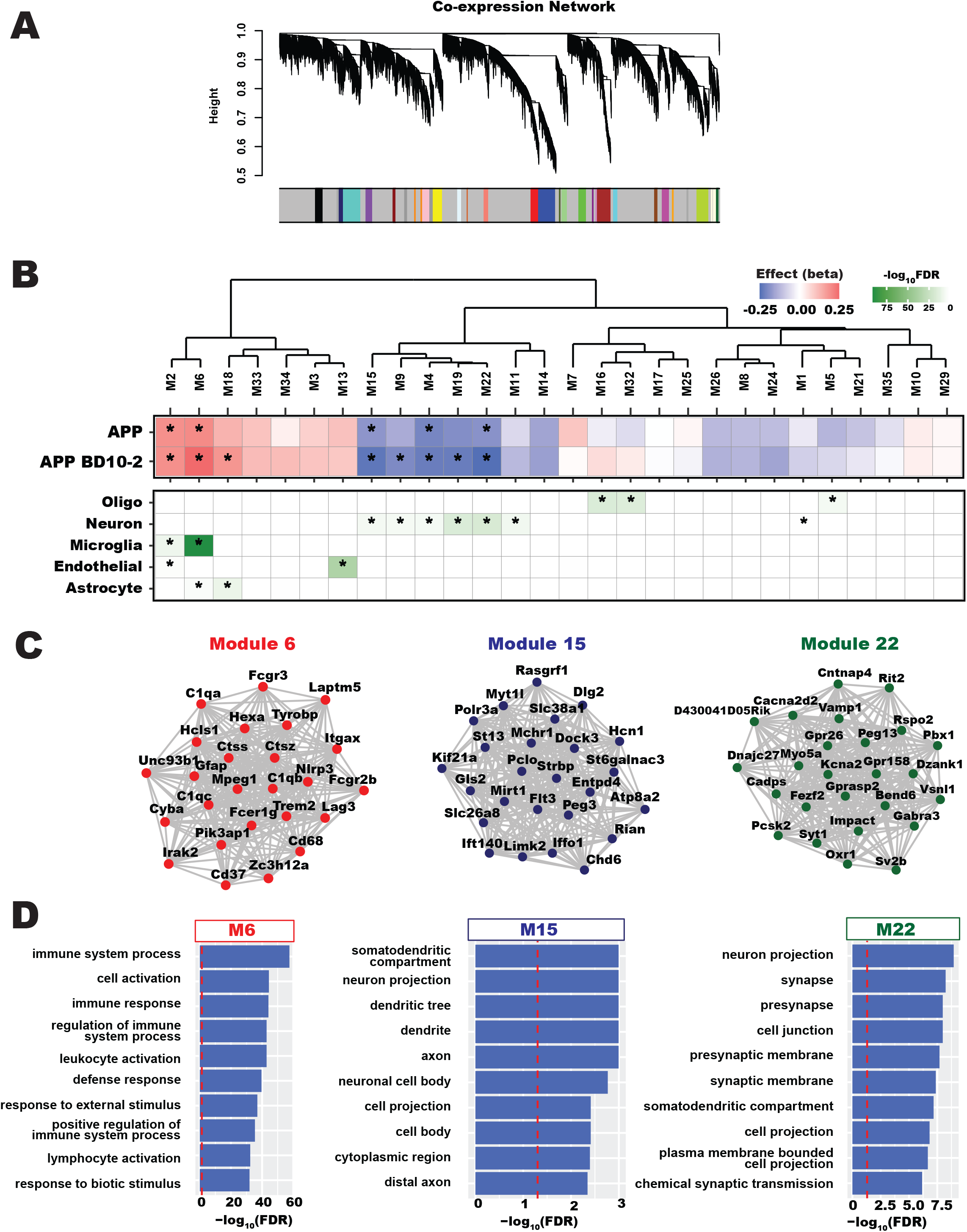
Weighted gene co-expression network analysis (WGCNA) of post-TBS hippocampal slices after 3 months in vivo BD10-2 treatment for 16-month-old APP^L/S^ mice. **A.** Hierarchical cluster tree of 23,407 expressed genes in stimulated hippocampal slices from WT-Veh mice and APP^L/S^ mice with or without BD10-2 treatment. The branches and color bands represent the 34 modules that were detected. **B.** Hierarchical clustering of gene co-expression modules by module eigengene in the APP effect (APP-Veh-TBS vs WT-Veh-TBS) and the APP-BD10-2 effect (APP-BD10-2-TBS vs WT-Veh-TBS). Only modules with a minimum size of 50 genes are shown. The plot below the dendrogram shows the module association strength with each effect. FDR < 0.05 for each module is denoted with an asterisk. A total of 5 modules were significantly associated with the APP effect. Three additional modules were significantly associated with the APP-BD10-2 effect. Module cell-type enrichments (*FDR < 0.05) are shown in the panel below for major central nervous system (CNS) cell types. **C.** Examples of specific modules dysregulated across APP and APP-BD10-2 effects, with the top 25 hub genes shown. **D.** Top 10 gene ontology (GO) enrichments for genes in modules 6, 15 and 22, red dashed line indicates FDR < 0.05 significance.

The five strongest downregulated co-expression modules (M15, M9, M4, M19, and M22) exhibited APP genotype associated downregulation and even stronger APP-BD10-2 associated downregulation (blue in upper panel of **Fig. 7B**), with two modules passing FDR significance (FDR < 0.05) for the APP-BD10-2 effect but not for the APP effect. These modules had significant cell type-specific enrichment for neurons (lower panel in **Fig. 7B**), concurring with the enrichment in neuronal AD-related co-expression modules (**Fig. 6E**). Modules M15 and M22 showed significant enrichment for dendrite, membrane projection and synaptic genes (**Fig. 7C, D**). Importantly, we also observed downregulation of synaptic genes and neuronal modules in the TBS effects from **Fig. 5**, suggesting again that this is a hallmark of BD10-2 mediated LTP rescue in APP^L/S^ mice. Finally, modules M32 and M16 showed significant enrichment of the oligodendrocyte cell type and appeared to have a slight opposing direction of effect in APP and APP-BD10-2 mice, but this was not FDR significant (**Fig. 7B**).

### 3.7 Transcription of genes involved in synaptic plasticity and immune related pathways was modified in stimulated hippocampal slices from BD10-2 treated APP^L/S^ mice

Given the very notable rescue of LTP deficits in APP^L/S^ mice when treated with BD10-2 and its action via TrkB/C signaling, we further examined functional pathways downstream of neurotrophin signaling that potentially impact synaptic plasticity. Looking at functional ontologies broadly, Milind et al [90] iteratively re-clustered the Wan et al. AD co-expression modules to develop pathway specific modules that we used for AD-related overlap enrichment (**Fig. 8A, Figure S4C**). Overlap enrichment for APP and APP-BD10-2 effect DE (padj < 0.05) in these modules show similar patterns of immune response, synaptic signaling, as well as enrichment of upregulated oligodendroglial & cytoskeletal rearrangement modules. This agrees with the overlap enrichment of Wan et al. modules (**Fig. 6E**) and co-expression module analysis (**Fig. 7**). Unsurprisingly, this carries forward into known LTP associated spine turnover ontologies. There is a broad enrichment of neurogenesis, cell membrane projection, combined with increased complement pathway and apoptotic processes, suggestive of considerable cellular turnover and endocytic activity **(Fig. 8B**). Many of these pathways are enriched in DE APP effect genes and are further enriched by the APP-BD10-2 effect (**Fig. 8B**). Overall, these enrichments again suggest that rather than counteracting aberrant gene expression in APP^L/S^ mice, BD10-2 acts on APP^L/S^ mice via potential compensatory mechanisms to enhance candidate beneficial pathways involved in response to the disease model.

**Figure 8.**
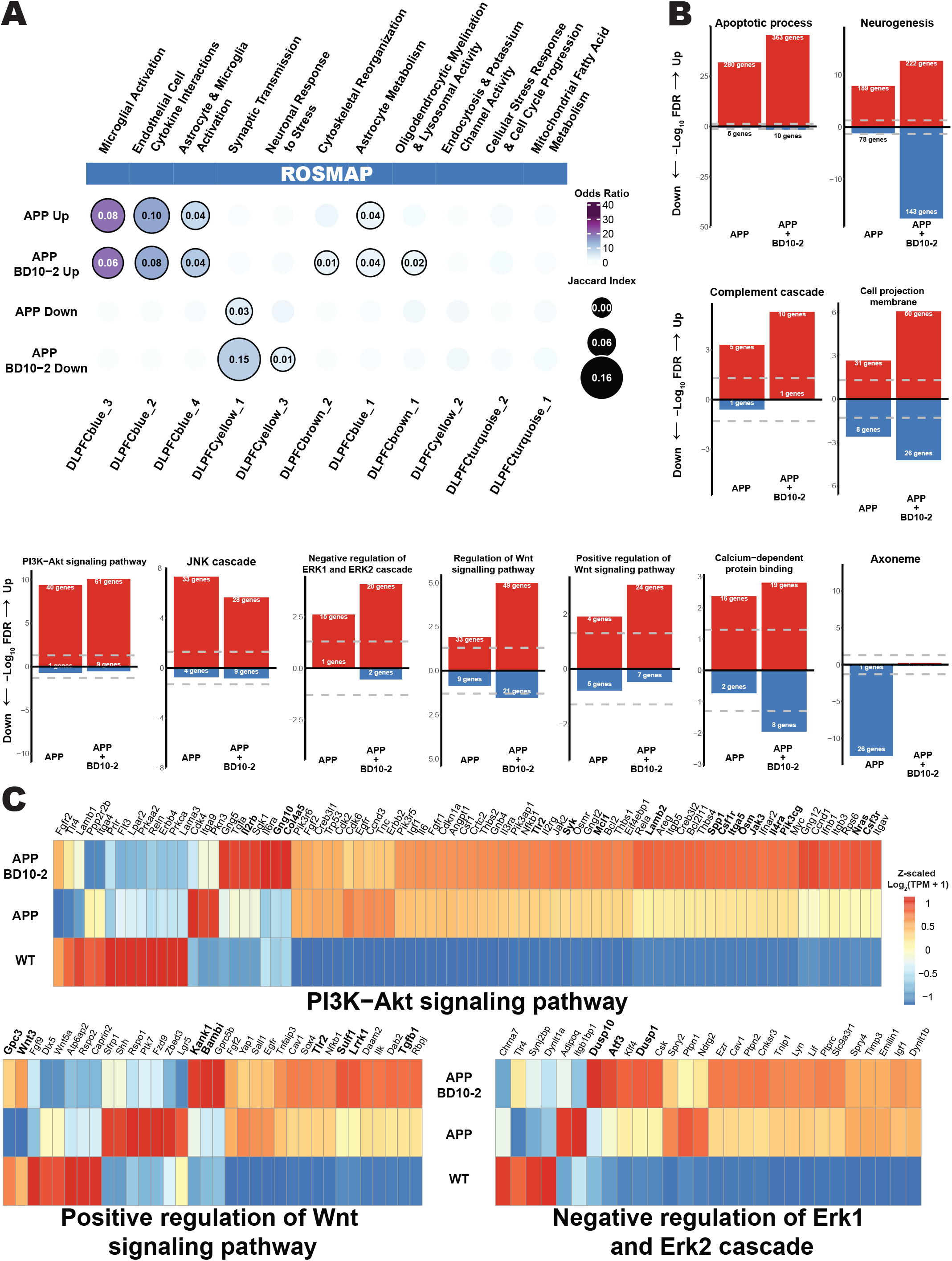
Pathway enrichment analysis of stimulated hippocampal slices from APP^L/S^ mice with or without BD10-2 treatment versus WT-Veh mice. **A.** Overlap enrichment (Fisher’s exact) of AD-related human-mouse co-expression modules from Milind et al. [90] compared with upregulated (top 2 rows)/downregulated (bottom 2 rows) DE genes (padj < 0.05) in the APP effect (APP-Veh-TBS vs WT-Veh-TBS) and the APP-BD10-2 effect (APP-BD10-2-TBS vs WT-Veh-TBS). Circles are sized and annotated with the Jaccard index of their overlap and are colored by the odds ratio of the overlap enrichment (circles with an FDR < 0.05 have a bold edge). **B.** Selected pathway enrichment in DE genes seen in the APP effect and the APP-BD10-2 effect. **C.** Examination of gene-scaled expression for genes associated with positive regulation of the Wnt signaling pathway (GO:0030177), PI3K-Akt signaling pathway (KEGG:04151), and negative regulation of ERK1 and ERK2 cascade (GO:0070373). Mean values for each group are Log_2_(TPM + 1), then z-scaled across samples. Gene names with nominal significance (p-value < 0.05) in BD10-2 effect (APP-BD10-2-TBS vs APP-Veh-TBS) stimulated slices are bolded.

Wnt, Erk, and PI3K-Akt pathways are downstream of TrkB and TrkC signaling and are thought to be candidate mechanisms for neurotrophin modulation of synaptic plasticity. The Wnt pathway showed enrichment in the APP effect with further enrichment in the APP-BD10-2 effect which incorporated a number of additional pathway genes that showed at least nominal significance in analysis of individual DE genes in the BD10-2 effect, i.e. Kank1, a modulator of apoptotic mechanisms and axonal function (**Fig. 6A, 8B, C, Table S9**) [91, 92]. The PI3K/Akt pathway has a known role in cell growth, neuronal development, and synapse formation (Sánchez-Alegría et al., 2018). While the extent of upregulation of PI3K/Akt pathway genes in the APP-BD10-2 effect was modest compared to the already observed increase in the APP effect, the total number of upregulated genes increased from 40 in the APP effect to 61 in the APP-BD10-2 effect. Additionally, negative regulation of Erk signaling, apoptotic processes, and complement cascade, were all enriched for upregulated genes (**Fig. 8B**). The terms neurogenesis, cell projection membrane, and calcium dependent protein binding’ were all enriched for both up and downregulated genes in APP, and APP-BD10-2 effects. The JNK cascade term showed a mild decrease in enrichment of upregulated genes (**Fig. 8B**). Looking at normalized expression of these pathways (**Fig. 8C**), the trend for many genes and pathways fits a potential compensatory pattern, with enrichment in APP^L/S^ mice and further enrichment with the addition of BD10-2. However, on a more granular level, there are genes that individually are differentially expressed in APP^L/S^ mice but rescued to match WT levels by BD10-2 treatment (**Fig. 8C**).

## 4 DISCUSSION

In the present study, we addressed the question of whether upregulation of TrkB/C receptor signaling, as achieved by administration of BD10-2, affects deficits in synaptic function present in APP^L/S^ mice. In vitro studies determined that BD10-2 promotes hippocampal neuron survival to a similar extent as its parent compound LM22B-10. Hippocampal slice and in vivo studies demonstrated that targeting of TrkB/C with BD10-2 ameliorates LTP and memory deficits present in APP^L/S^ mice in association with activating AKT and CaMKII signaling and restoring altered AMPAR subunit (GluA1) phosphorylation and localization, which are crucial to LTP. We also show that BD10-2 rescues activity-dependent, LTP-associated synaptic gene transcription and broadly activates transcription associated with microglial and immune pathways.

### APP and BD10-2 effects on signaling in late phase LTP

We first demonstrated that BD10-2, like LM22B-10, promotes cell survival acting through TrkB and TrkC receptors. It is well known that interaction of TrkB with its endogenous ligand BDNF modulates synaptic transmission and induces LTP [65, 93–96]. Consistent with this effect of BDNF, we found that BD10-2 can prevent synaptic plasticity and memory deficits that occur in aged APP^L/S^ mice suggesting that BD10-2 mimics, in part, the biological effects of BDNF by targeting TrkB/C. We found that major components of synaptic plasticity are restored in APP^L/S^ mice by targeting TrkB/C receptors in vivo, even though the compound was not physically present at the time of slice recording, suggesting that BD10-2 might cause persistent structural and functional changes at synapses that contribute to recovering their function.

Initiation of hippocampal LTP requires calcium-dependent activation of CaMKII at dendritic spines which promotes the activation, trafficking, and surface insertion of synaptic AMPARs [97–103]. Later stages of LTP require changes in gene expression and protein synthesis and establish long lasting changes to the synapse that are maintained over time through persistent biochemical changes in the cell [65, 104, 105]. LTP activates phosphorylation of many proteins such as CaMKII, GluA1-containing AMPARs, as well as Akt, and ERK, which are crucial for inducing and maintaining synaptic plasticity through regulation of activity-induced protein synthesis [68, 97, 106–111]. Aberrant regulation of these proteins has been reported in AD human brain and several AD mouse models [112, 113]. BDNF-TrkB signaling activates many of these same pathways including the canonical three: PI3K/AKT, MAPK/ERK, and PLCγ/PKC pathways and is directly involved in the maintenance of late phase LTP [65]. Notably, activation of the PLCγ/PKC pathway can trigger Ca^2+^ influx which can activate CaMKII which is involved in LTP induction [62–66]. Signaling through PLCγ/PKC, MAPK/ERK, and PI3K/AKT pathways can all induce downstream activation of transcription factors such as cyclic-AMP responsive element binding protein (CREB) which promotes expression of genes required for LTP maintenance [65, 114–116]. We examined the phosphorylation as an indicator of activity and levels of proteins involved in these three pathways (PI3K/AKT, MAPK/ERK, and PLCγ/PKC). Our data, consistent with Lee et al (2010), indicates a significant decrease in the phosphorylation of the AMPAR subunit GluA1 and its kinases CAMKII(-α and -β) as well as altered activity-dependent colocalization of GluA1 with the dendritic protein drebrin in APP^L/S^ mice. Phosphorylation of GluA1 and CaMKII-β (but not CaMKII-α) and as well as GluR1/drebrin colocalization was preserved at WT levels by chronic BD10-2 treatment. Drebrin is an actin-binding protein that promotes dendritic growth at excitatory post synapses [117, 118]. AMPAR activity stabilizes drebrin in spines [119]. Our data suggests that modulation of TrkB/C receptors by BD10-2 treatment contributes to AMPAR-dependent stabilization of drebrin in spines as a novel activity-dependent mechanism for normalization of APP/Aβ-associated synaptic dysfunction. Remarkably, phosphorylation of CaMKII-β, and AKT were decreased and restored by BD10-2 only in non-stimulated slices from APP^L/S^ mice while deficits seen in GluA1 phosphorylation in APP^L/S^ mice were only restored by BD10-2 in stimulated slices indicating that rescue of GluA1 activation by BD10-2 is activity-dependent. Perhaps rescue of baseline levels of these proteins is sufficient to restore downstream activity-dependent mechanisms such as synaptic recruitment of drebrin and activated AMPARs. Based on these findings, we propose a model, similar to what has been described previously for BDNF [65], in which BD10-2 prevents APP/Aβ-induced deficits in early phases of LTP by promoting CaMKII and AMPA receptor phosphorylation/activation through activation of the TrkB/C-regulated PLCγ/PKC pathway, thereby rescuing activity dependent AMPAR/drebrin colocalization in dendritic spines.

### Effects of APP and BD10-2 on activity-dependent late phase LTP associated transcription

We next asked whether APP^L/S^ expression (described here as the APP effect) and/or BD10-2 influenced activity-dependent LTP associated gene expression which is required to support protein synthesis alterations needed for long lasting synaptic changes occurring in late-phase LTP. RNA-seq on 130 min (late-phase) post-TBS hippocampal brain slices revealed LTP dependent downregulation of genes involved in synaptic signaling and potentiation. Earlier timepoints would be needed to determine if there is a post-TBS timepoint at which these synaptic genes are upregulated; nevertheless, there is a clear transcriptomic signature of LTP activity that is not found in slices that did not undergo LTP (TBS-): occurring in WT-Veh mice, altered in APP-Veh mice, and restored in APP-BD10-2 mice. Overlap enrichment analysis revealed that these transcriptional changes overlap with neuronal modules previously determined to have relevance to AD. Direct comparison of TBS+ slices for the APP effect and the APP-BD10-2 effect also showed that this signature was present only with the addition of BD10-2, further implicating the compound in LTP rescue in an AD context. Together with direct observation of LTP on these same hippocampal slices we see a measurable and lasting transcriptomic impact during the rescue of LTP by BD10-2.

### Transcriptional effects of APP and BD10-2 on microglial genes in the context of LTP

We observed strong upregulation of broad sets of immune relevant genes and pathways with TBS stimulation regardless of whether the samples had measurable LTP after TBS. Compared to WT, these pathways exhibited stronger upregulation in APP^L/S^ mice with and without BD10-2 suggesting this immune upregulation is not associated with activity-dependent LTP but may be correlated with amyloid pathology which is not the direct target of BD10-2. Overlap enrichment analysis and network module analysis of the APP-effect in stimulated samples reiterated this broad upregulation of AD relevant genes but only mildly increases with the addition of BD10-2. However, within the immune & microglial enrichments, microglia-associated pathways—including the complement pathway—were more enriched in response to BD10-2 and stimulation.

Notable genes that were upregulated in both the APP effect and further upregulated in the APP-BD10-2 effect include Clec7a and Cst7, two markers of disease associated microglia [120–122] (**Fig. 6C**). Our data also identifies a gene transcription module (M6red), which is enriched for a large set of complement genes that have previously been linked to microglial response to Aβ: Trem2, Tyrobp, Cd68, C1qa, C1qb, C1qc, and C3or. Microglia are the dominant immune cell type in the brain [123] and they play a major role in Aβ clearance [124]. Moving beyond amyloid, one of the top DE genes in the BD10-2 effect itself was C3, still significantly upregulated over 24 hours after clearance of BD10-2 from the brain, suggesting a lasting engagement of this specific complement mechanism (see also **Fig. 8B**). C3 has been shown to label synapses targeted for synaptic pruning, involving the complement system directly in synapse elimination as a result of LTP promoted spine turnover [125, 126]. Alterations in the complement system have been observed in AD patients [127] and AD mice [88], and excessive synaptic elimination in AD suggests the complement system balances microglial response between protective and detrimental [128–131]. Upregulation of these complement genes was correlated with an enrichment of neurogenesis, apoptosis, and neurotrophin pathways impacting synaptic plasticity/turnover, but further studies will be needed to understand the mechanisms by which LTP may drive this balance.

### Limitations and conclusions

Targeting TrkB/C receptor activation is a promising candidate for translational drug development particularly because it rescues synaptic function and cognition in mice with late-stage pathology. Pathological changes may accumulate for years prior to clinical detection of AD biomarkers or symptom onset. We found that targeting of TrkB/C receptors with BD10-2, rescues LTP and cognition, modulates LTP associated pathways downstream of TrkB signaling, and alters activity-dependent gene expression. To assess the large volume of data generated in our RNA-seq studies of transcriptional changes that occur in L-LTP, we focused on analyzing broad changes in related genes using various computational methods, however, it is possible we are missing more granular changes that occur in specific genes or pathways. Our data generate new questions surrounding activity dependent transcription of synaptic and immune programs during LTP that will need to be explored in future studies, especially in the context of AD pathophysiology and therapeutic target development.

## Data Availability

The raw data for this study has been deposited in the NCBI’s Gene Expression Omnibus under GSE243110. Interactive dashboards of study data are available at the project website: https://longo-lab.github.io/docs/projects/T41B_BD10-2_stim.html. The consensus clusters used for the Wan et al co-expression analysis are available at https://doi.org/10.7303/syn10309369.1.

## Code Availability

Where possible, code for generating the raw results is available at the project website: https://longo-lab.github.io/docs/projects/T41B_BD10-2_stim.html. Code for the processing of RNAseq data using the encode long-rnas pipeline is available at https://www.encodeproject.org/data-standards/rna-seq/long-rnas/. Any code not available via the website can be provided by request to the corresponding author.

## Acknowledgements

We would like to thank the Stanford Functional Genomics Facility for performing the RNA-seq experiments. Some of the computing for this project was performed on the Stanford SCG cluster. We would like to thank the Stanford Research Computing Center for providing computational resources and support. Wan et al network modules were based on data obtained from the AD Knowledge Portal. Data generation was supported by the following NIH grants: P30AG10161, P30AG72975, R01AG15819, R01AG17917, R01AG036836, U01AG46152, U01AG61356, U01AG046139, P50 AG016574, R01 AG032990, U01AG046139, R01AG018023, U01AG006576, U01AG006786, R01AG025711, R01AG017216, R01AG003949, R01NS080820, U24NS072026, P30AG19610, U01AG046170, RF1AG057440, and U24AG061340, and the Cure PSP, Mayo and Michael J Fox foundations, Arizona Department of Health Services and the Arizona Biomedical Research Commission. We thank the participants of the Religious Order Study and Memory and Aging projects for the generous donation, the Sun Health Research Institute Brain and Body Donation Program, the Mayo Clinic Brain Bank, and the Mount Sinai/JJ Peters VA Medical Center NIH Brain and Tissue Repository. Data and analysis contributing investigators include Nilüfer Ertekin-Taner, Steven Younkin (Mayo Clinic, Jacksonville, FL), Todd Golde (University of Florida), Nathan Price (Institute for Systems Biology), David Bennett, Christopher Gaiteri (Rush University), Philip De Jager (Columbia University), Bin Zhang, Eric Schadt, Michelle Ehrlich, Vahram Haroutunian, Sam Gandy (Icahn School of Medicine at Mount Sinai), Koichi Iijima (National Center for Geriatrics and Gerontology, Japan), Scott Noggle (New York Stem Cell Foundation), Lara Mangravite (Sage Bionetworks). We express our special gratitude to Tariq Ahmed for his profound intellectual contributions and innovative scientific insights, which were instrumental in deciphering the synaptic mechanisms proposed in this paper. Additionally, we appreciate his invaluable supervision and guidance during the execution of electrophysiology experiments, which greatly advanced our research.

## Author contributions

A.L.H. and T.Y. designed and performed experiments and analyzed the data. P.S.M. performed behavioral experiments and analyzed the data. R.R.B., P.M.L and T.W.C. analyzed the RNA-seq data. H.W. analyzed immunostaining data. K.C.T and H.L administered the compound to the four cohorts of animals. A.L.H., T.Y., R.R.B, V.F.L. and D.A.S wrote the manuscript. A.L.H and F.M.L conceived and supervised the project, designed the study and interpreted the data, and finalized the manuscript.

