## Supplementary figures, tables and legends for "A TrkB and TrkC partial agonist restores deficits in synaptic function and promotes activity-dependent synaptic and microglial transcriptomic changes in a late-stage Alzheimer’s mouse model"

**Legends to supplementary figures and tables:**

Supplementary Fig. 1**. Effect of BD10-2 on proteins downstream of TrkB signaling in WT and APP$^{L/S}$ mice.** BD10-2 or vehicle was administered by oral gavage to 13-month-old APP$^{L/S}$ mice for three months, followed by collection of hippocampal slices. Western blots of hippocampal slice extracts were quantitated by determining the ratios of phospho (p) protein over total protein or total protein over actin, and then normalized to respective WT-Veh groups. Statistical significance was determined using either one way ANOVA with Sidak post hoc test or Kruskal-Wallis test with post-hoc Dunn's multiple comparisons test (mean ± SEM, Sample size = 6-10 hippocampal slices, 3-5 mice per group, with two independent western blots averaged per slice). **A-E** Quantification of TBS- hippocampal slice blots shown in **Fig. 4A**. **F.** Representative western blots for each TBS+ condition, except for GluA1 which is shown in **Fig. 4I**. **G-Q.** Quantification of TBS+ hippocampal slices shown in **Fig S1F** and **Fig 4I**.

Supplementary Fig. 2. **Transcriptomic sample quality control and outlier detection**. **(A)** Heatmap of sample-to-sample Poisson distances clustered hierarchically. **(B)** Standardized sample network connectivity colored by each group. Dashed lines at |Z score| > 2.

Supplementary Fig. 3. **Volcano plots of differential expression (DE) for TBS, APP BD10-2, and APP-BD10-2 effects**. Horizontal lines represent nominal significance of p-value < 0.05, vertical lines indicate fold changes of ± $\log_2(1.2)$. **(A)** Volcano plots examining the TBS effect in experimental groups. *Left:* the TBS effect in WT-Veh mice (WT-Veh-TBS vs WT-Veh); *Center:* the TBS effect in APP-Veh mice (APP-Veh-TBS vs APP-Veh); *Right:* the TBS effect in APP-BD10-2 mice (APP-BD10-2-TBS vs APP-BD10-2). **(B)** Volcano plots of differential expression in stimulated hippocampal slices. *Left:* the APP effect (APP-Veh-TBS vs WT-Veh-TBS); *Center:* the APP-BD10-2 effect (APP-BD10-2-TBS vs WT-Veh-TBS); *Right:* the BD10-2 effect (APP-BD10-2-TBS vs APP-Veh-TBS).

Supplementary Fig. 4. **Full overlap enrichment analysis of differential expression datasets**. **(A)** Overlap enrichment (Fisher's exact) of AD-related human-mouse co-expression modules from Wan et al. [52] compared with upregulated (top 3 rows)/downregulated (bottom 3 rows) genes (padj < 0.05) in the TBS effect in the three treatment groups: WT-Veh (WT-Veh-TBS vs WT-Veh), APP-Veh (APP-Veh-TBS vs APP-Veh), and APP-BD10-2 (APP-BD10-2-TBS vs APP-BD10-2). Circles are sized and annotated with the Jaccard index of their overlap and are colored by the significance of the overlap enrichment (-log FDR; circles with an FDR < 0.05 have a bold edge). Modules are grouped based on "consensus clusters", yellow, Astroglial-like modules; light blue, Microglial-like modules; maroon, Neuronal-associated modules; green, Oligodendroglial-like modules. Modules are additionally annotated along the bottom by the associated brain region of the human AD cohort from which they were derived, CBE, cerebellum; DLPFC, dorsolateral prefrontal cortex; FP, frontal pole; IFG, inferior frontal gyrus; PHG, parahippocampal gyrus; STG, superior temporal gyrus; TCX, temporal cortex. **(B)** Overlap enrichment (Fisher's exact) of AD-related human-mouse co-expression modules from Wan et al. [52] compared with upregulated (top 3 rows)/downregulated (bottom 3 rows) DE genes (padj < 0.05) in the APP effect (APP-Veh-TBS vs WT-Veh-TBS), the BD10-2 effect (APP-BD10-2-TBS vs APP-Veh-TBS), and the APP-BD10-2 effect (APP-BD10-2-TBS vs WT-Veh-TBS). **(C)** Overlap enrichment (Fisher's exact) of AD-related human-mouse co-expression modules from Milind et al. [90] compared with upregulated (top 2 rows)/downregulated (bottom 2 rows) DE genes (padj < 0.05) in the APP effect and the APP-BD10-2 effect. Circles are sized and annotated with the Jaccard index of their overlap and are colored by the odds ratio of the overlap enrichment (circles with an FDR < 0.05 have a bold edge).

Supplementary Table S1. **Differential Gene Expression of WT VEH TBS vs WT VEH.**

Supplementary Table S2. **Differential Gene Expression of APP VEH TBS vs APP VEH.**

Supplementary Table S3. **Differential Gene Expression of APP BD10-2 TBS vs APP BD10-2.**

Supplementary Table S4. **Gprofiler Pathway Enrichment of WT VEH TBS vs WT VEH.**

Supplementary Table S5. **Gprofiler Pathway Enrichment of APP VEH TBS vs APP VEH.**

Supplementary Table S6. **Gprofiler Pathway Enrichment of APP BD10-2 TBS vs APP BD10-2.**

Supplementary Table S7. **Gprofiler Pathway Enrichment of Quadrants of Figure 5B (Comparing TBS effect in WT and APP$^{L/S}$ mice).**

Supplementary Table S8. **Differential Gene Expression of APP VEH TBS vs WT VEH TBS.**

Supplementary Table S9. **Differential Gene Expression of APP BD10-2 TBS vs APP VEH TBS.**

Supplementary Table S10. **Differential Gene Expression of APP BD10-2 TBS vs WT VEH TBS.**

Supplementary Table S11. **Gprofiler Pathway Enrichment of APP VEH TBS vs WT VEH TBS.**

Supplementary Table S12. **Gprofiler Pathway Enrichment of APP BD10-2 TBS vs APP VEH TBS.**

Supplementary Table S13. **Gprofiler Pathway Enrichment of APP BD10-2 TBS vs WT VEH TBS.**

Supplementary Table S14. **Gprofiler Pathway Enrichment of Quadrants of Figure 6C (Comparing APP and BD10-2 Expression).**

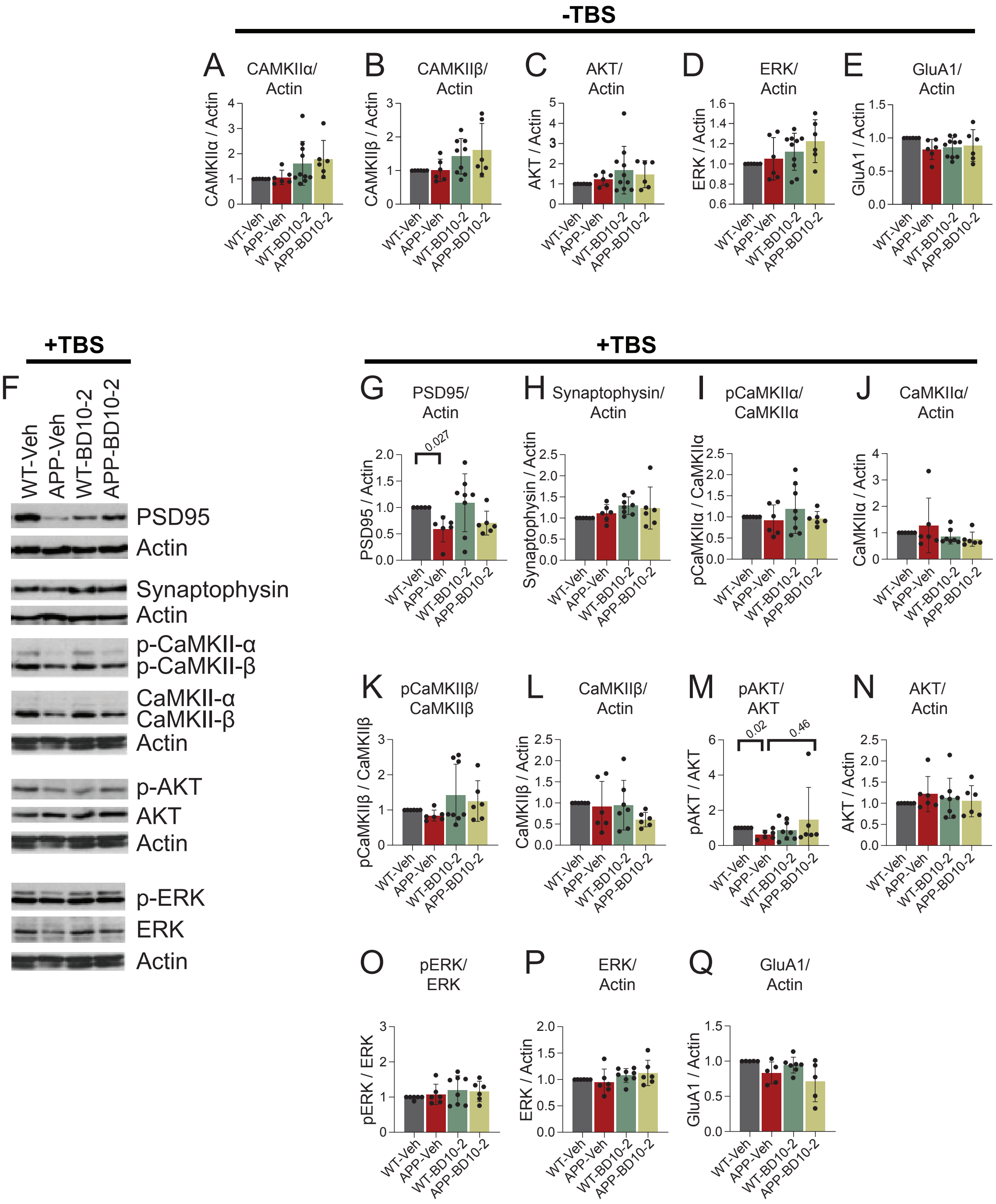


### Supplementary Figure 1

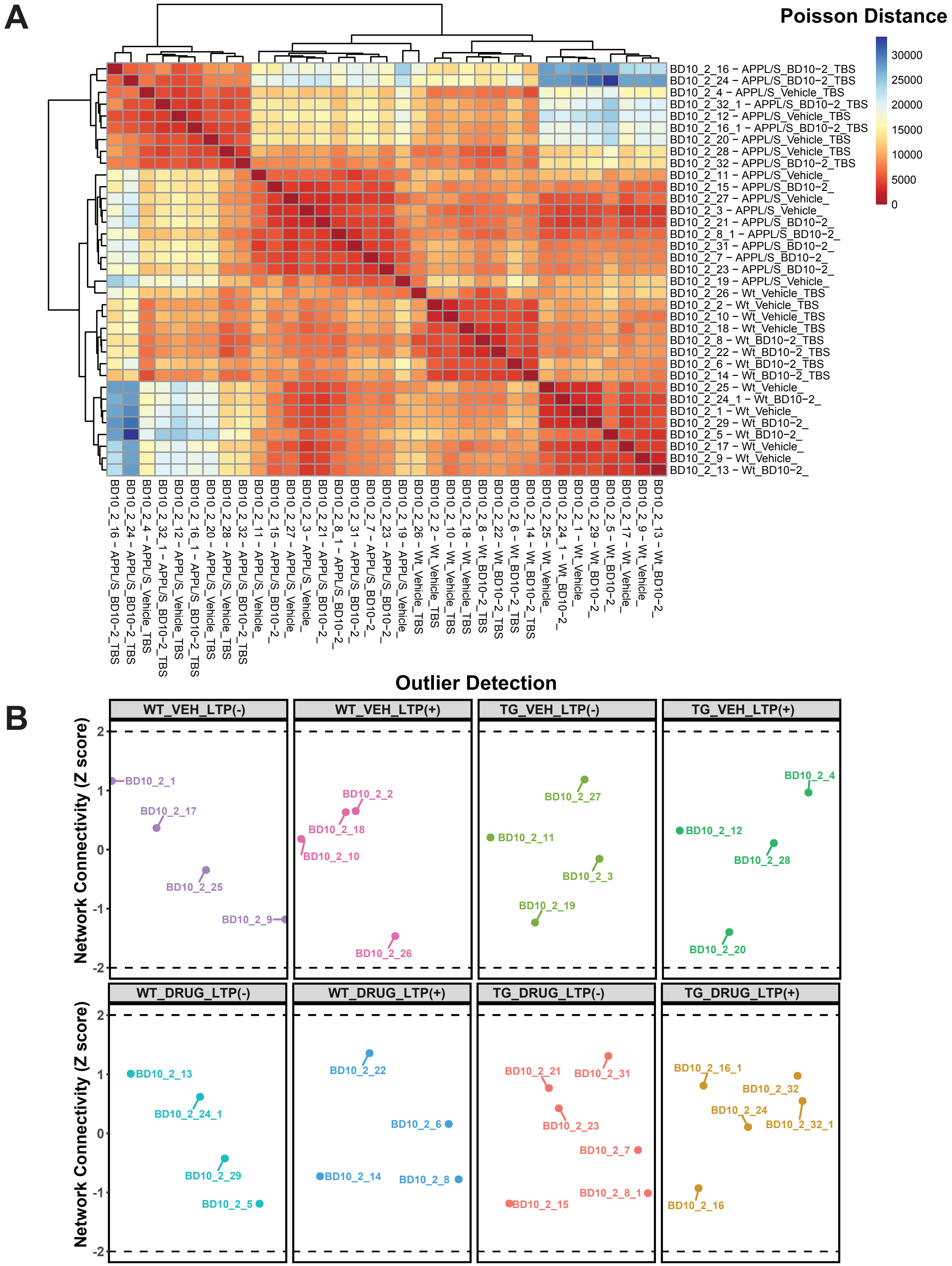


### Supplementary Figure 2

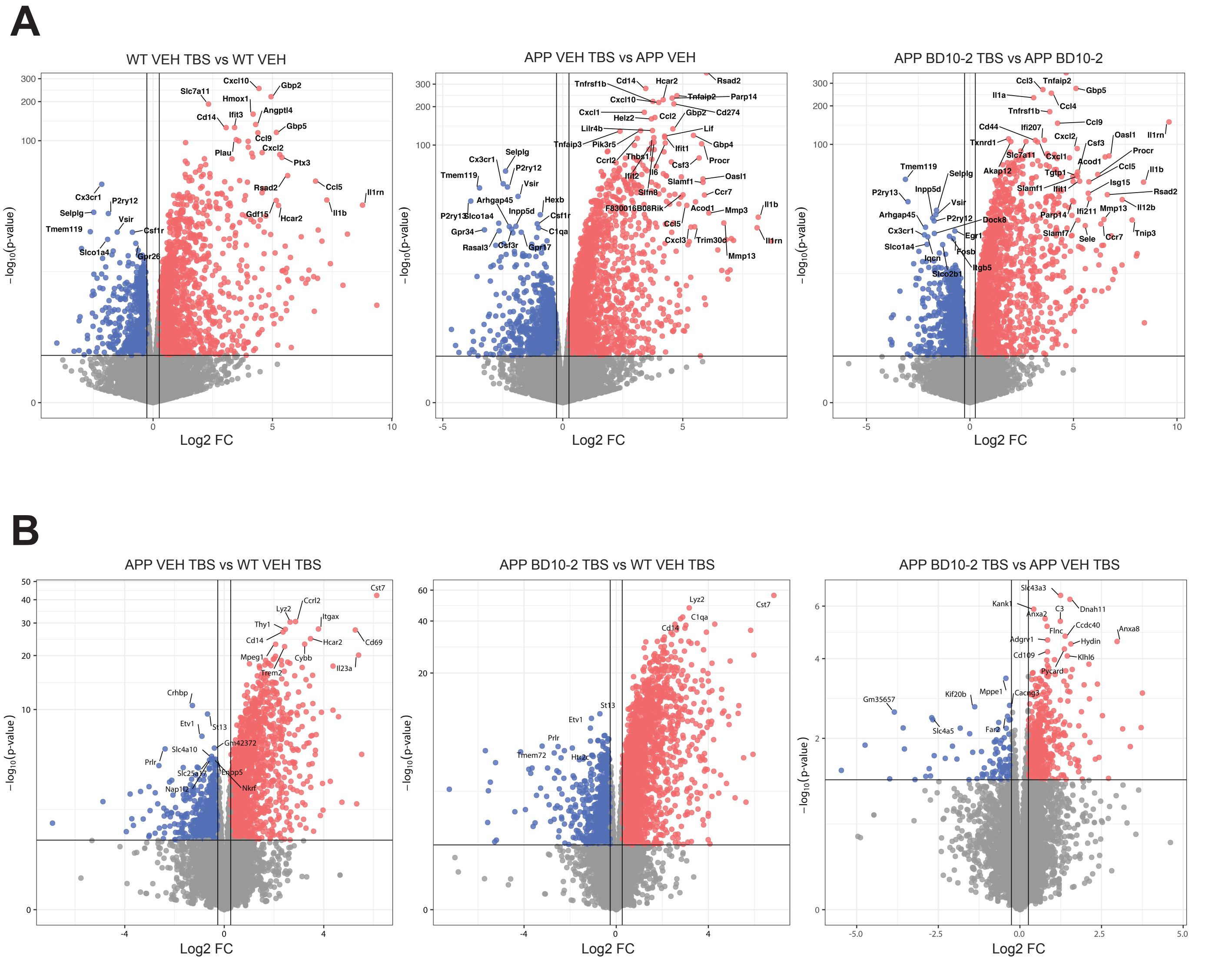


### Supplementary Figure 3

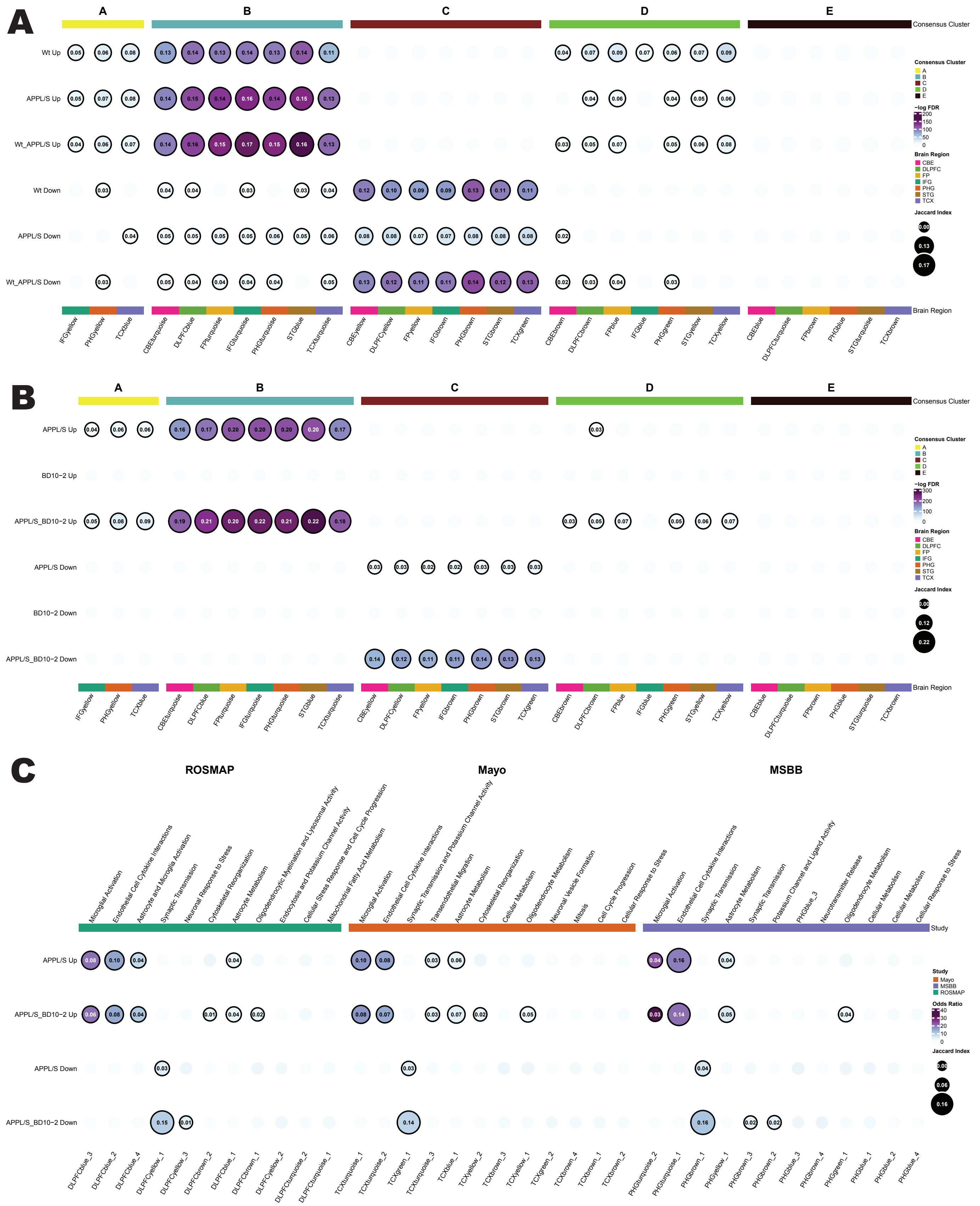


### Supplementary Figure 4
